# AGAMOUS mediates timing of guard cell formation during gynoecium development

**DOI:** 10.1101/2023.01.23.525231

**Authors:** Ailbhe J. Brazel, Róisín Fattorini, Jesse McCarthy, Rainer Franzen, Florian Rümpler, George Coupland, Diarmuid S. Ó’Maoiléidigh

## Abstract

In *Arabidopsis thaliana*, stomata are composed of two guard cells that control the aperture of a central pore to facilitate gas exchange between the plant and its environment, which is particularly important during photosynthesis. Although leaves are the primary photosynthetic organs of higher plants, floral organs are also photosynthetically active. In the Brassicaceae, evidence suggests that silique photosynthesis is important for optimal seed oil content. A group of transcription factors containing MADS DNA binding domains is necessary and sufficient to confer floral organ identity. Elegant models, such as the ABCE model of flower development and the floral quartet model, have been instrumental in describing the molecular mechanisms by which these floral organ identity proteins govern flower development. However, we lack a complete understanding of how the floral organ identity genes interact with the underlying leaf development program. Here, we show that the MADS domain transcription factor AGAMOUS (AG) represses stomatal development on the gynoecial valves, so that maturation of stomatal complexes coincides with fertilization. We present evidence that this regulation by AG is mediated by direct transcriptional repression of the master regulator of the stomatal lineage, *MUTE*, and that this interaction is conserved among the Brassicaceae. This work extends on our understanding of the mechanisms underlying floral organ formation and provides a framework to decipher the mechanisms that control floral organ photosynthesis.

## Introduction

In eudicots, such as *Arabidopsis thaliana*, flowers are composed of four types of floral organs: sepals, petals, stamens, and carpels. The sepals contain high levels of chlorophyll and bear stomata, making them leaf-like in appearance. In contrast, mature petals and stamens lack substantial concentrations of chlorophyll (Smyth *et al*., 1990; Pyke and Page, 1998). Stomata are absent from petals but modified stomata are present on the abaxial surfaces of anthers (Smyth *et al*., 1990; Nadeau and Sack, 2002*a*). In *A. thaliana*, the ovules are encased within the gynoecium which is formed of two fused carpels and other tissues that arise from the carpels, such as the style, stigma and replum (Smyth *et al*., 1990; Alvarez and Smyth, 2002). Although the gynoecial valves lack stomata, they are present on the differentiated and elongated silique valve epidermis (Geisler *et al*., 1998; Gu *et al*., 1998). Stomata on siliques enable atmospheric carbon fixation to support local photosynthesis and photosynthetic activity of siliques has been demonstrated in several members of the Brassicaceae family including *A. thaliana* (Sheoran *et al*., 1991; Gammelvind *et al*., 1996; Wang *et al*., 2016; Zhu *et al*., 2018). This photosynthetic activity positively influences seed oil content and is of interest to crop breeders (Hua *et al*., 2012; Wang *et al*., 2016; Zhu *et al*., 2018). Stomata also support transpiration, which drives the movement of nutrients through the plant and simultaneously facilitates cooling (Cramer *et al*., 2009). However, very little is known about the molecular mechanisms of stomatal development on these organs.

On leaves of *A. thaliana*, the stomatal cell lineage is initiated by the asymmetric cell division of a protodermal cell (or meristemoid mother cell), which is regulated by the basic helix-loop-helix (bHLH) transcription factor SPEECHLESS (SPCH) (MacAlister *et al*., 2007; Dong and Bergmann, 2010; Torii, 2021) (Supplemental Figure 1A). This asymmetric division produces a meristemoid and a stomatal lineage ground cell (SLGC). The meristemoid may then undergo further asymmetric cell divisions to renew itself and produce more SLGCs. Alternatively, a meristemoid may transition into a rounded cell termed a guard mother cell (GMC) (Dong and Bergmann, 2010; Torii, 2021). The transition from meristemoid to GMC is coordinated by another bHLH transcription factor, MUTE, which also functions to attenuate asymmetric cell divisions (Pillitteri *et al*., 2007). GMCs then divide symmetrically to produce two guards cells and ultimately a mature stomatal complex, a process that is mediated by a third bHLH transcription factor, FAMA (Ohashi-Ito and Bergmann, 2006). Two other bHLHs, SCREAM (SCRM) and SCRM2, form heterodimers with SPCH, MUTE, and FAMA to coordinate gene expression (Kanaoka *et al*., 2008). These division steps are regulated such that stomatal complexes are separated by at least one cell, which ensures control of the central pore aperture (Dong and Bergmann, 2010; Torii, 2021). Several transmembrane receptors have been implicated in this stomatal patterning, such as TOO MANY MOUTHS (TMM), ERECTA (ER), ER-LIKE1 (ERL1), and ERL2 (Nadeau and Sack, 2002*b*; Shpak *et al*., 2005; Torii, 2021). The activities of these receptors are modulated by the EPIDERMAL PATTERNING FACTORs (EPFs) and EPF-LIKES (EPFLs) secreted peptide families (Supplemental Figure 1A) (Hara *et al*., 2007; Hunt and Gray, 2009; Kondo *et al*., 2010; Torii, 2021).

**Figure 1.**
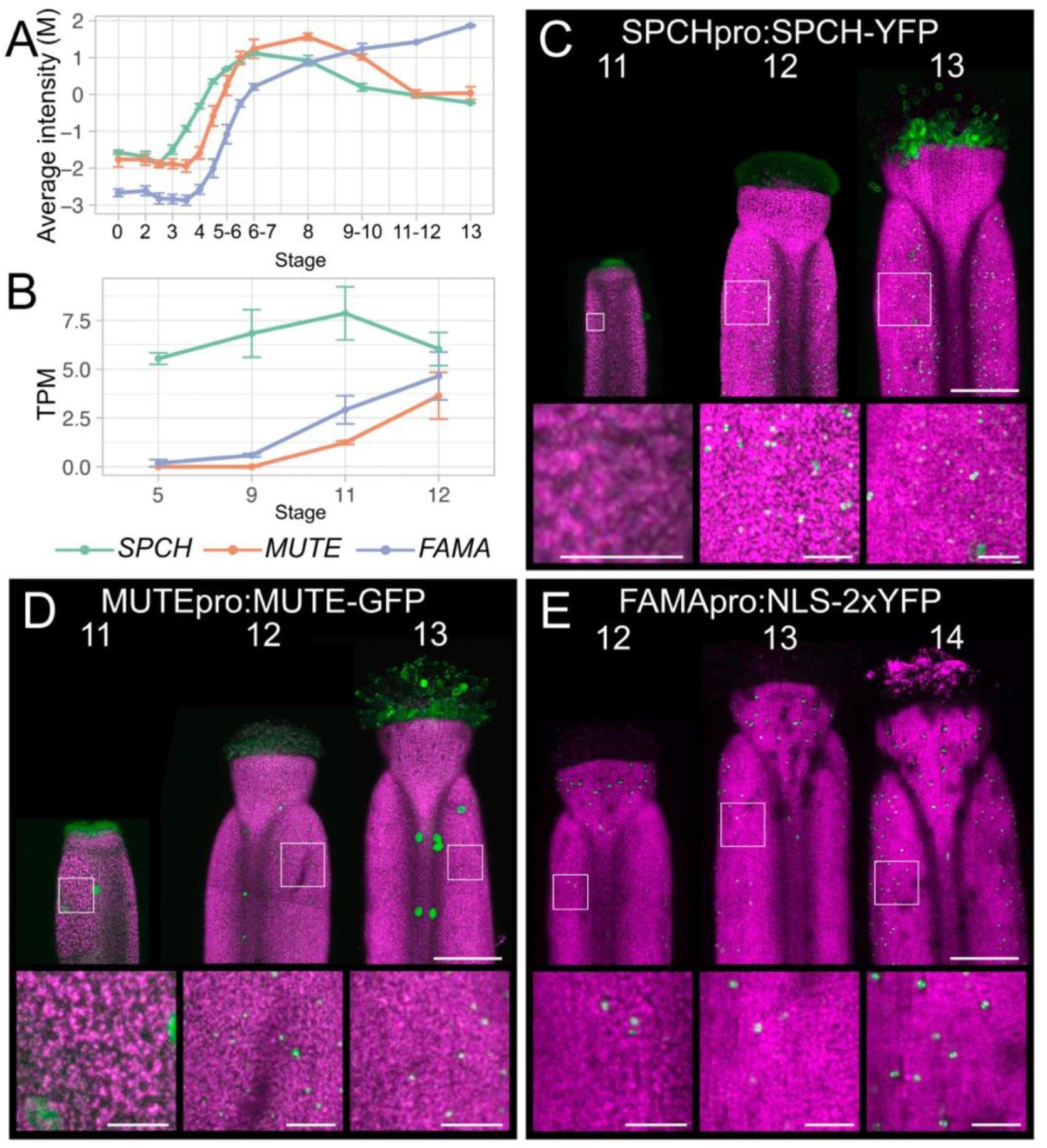
Expression of stomatal bHLH transcription factors during gynoecium development. (A-B) Levels of *SPCH, MUTE*, and *FAMA* mRNAs at described stages of flower development in (A) whole flower buds (Ryan *et al*. 2015) and (B) laser-microdissected gynoecia (Kivivirta *et al*. 2020). Error bars in (A-B) are s.e.m of three and four biological replicates, respectively. M, log-ratio of time-point to a common reference; TPM, transcripts per million. (C-E) Maximum intensity projections of stitched confocal laser scanning z-stacked micrographs of (C) *SPCHpro:SPCH-YFP*, (D) *MUTEpro:MUTE-GFP*, and (E) *FAMApro:NLS-2xYFP*, transgenes at different stages of gynoecium development as indicated. White boxes indicate areas that were magnified to produce the insets. Fluorescent protein is coloured green and chlorophyll fluorescence is coloured magenta. Scale bars for images of whole gynoecia are 100 µm. Scale bars for insets are 20 µm.

Floral organ identity is controlled by a group of transcription factors that contain MADS DNA-binding domains (Bowman *et al*., 1989, 1991*b*; O’Maoileidigh *et al*., 2014*a*; Sablowski, 2015; Thomson and Wellmer, 2019). In the absence of their activities, floral organs are converted into leaf-like organs while ectopic expression of these transcription factors is sufficient to transform leaves into floral organs (Bowman *et al*., 1991*b*; Pelaz *et al*., 2000*a*, 2001; Honma and Goto, 2001; Ditta *et al*., 2004). These observations confirmed a long-standing hypothesis that floral organs are derived from leaves (Goethe, 1790). They also formed the basis of the ABCE model of flower development and the floral quartet model, which largely address organ specification (Bowman *et al*., 1991*b*; Pelaz *et al*., 2000*b*, 2001; Honma and Goto, 2001; Ditta *et al*., 2004; Theißen *et al*., 2016). However, these MADS domain transcription factors continue to be expressed after the floral organs have been specified and they are known to control the expression of genes required for differentiation (Ito *et al*., 2004, 2007; O’Maoileidigh *et al*., 2014*b*; Ó’Maoiléidigh *et al*., 2018). Identifying the repertoire of differentiation processes that the floral organ identity genes control remains a key challenge (Sablowski, 2015).

The MADS-domain protein AGAMOUS (AG) controls the specification of stamens and carpels and is required for floral meristem termination (Bowman *et al*., 1989). AG interacts with E-class proteins, SEPALLATA 1-4 (SEP1-4), in heterodimeric or tetrameric complexes to coordinate carpel specification and floral meristem termination, with SEP3 playing an especially important role (Honma and Goto, 2001; Immink *et al*., 2009; Theißen *et al*., 2016). More recent data, however, suggests that the AG-SEP3 tetramers are only required for floral meristem termination rather than organogenesis (Hugouvieux *et al*., 2018). *AG* activity promotes carpel development in a partially redundant manner with its closest related paralogs *SHATTERPROOF1* (*SHP1*) and *SHP2* (Favaro *et al*., 2003). AG and the SHP proteins also suppress the formation of epidermal hairs (trichomes) during carpel differentiation, which represents the first tangible example of how the floral organ identity proteins modify the underlying leaf development program to generate floral organs (Ó’Maoiléidigh *et al*., 2013, 2018).

Here, we investigate the developmental progression and molecular regulation of stomatal development on gynoecial and silique valves in *A. thaliana*. We describe the normal progression of stomatal development before and after fertilization, and present evidence that AG suppresses this process in *A. thaliana* and other members of the Brassicaceae. The data presented provide further evidence and mechanism that transcription factors conferring floral organ identity directly suppress aspects of leaf development during floral organ formation. They also provide a framework with which to understand the establishment of the silique as a photosynthetic organ in the Brassicaceae.

## Materials and methods

### Plant growth and materials

Plants were grown on soil under cool white light at ~20°C in 16:8 h light:dark conditions. Plant lines used in this study are listed in Supplemental Table 1 Flower development stages are after Smyth *et al*., 1990.

### Genotyping

Genotyping PCRs were performed with genomic DNA extracted according to Edwards *et al*., 1991. Tissue was disrupted using a TissueLyser II (Qiagen). Primers used in this study are listed in Supplemental Table 2. The genotyping of *ag-10* and *ful-1* was performed as in Liu *et al*., 2011 and (Ferrándiz *et al*., 2000*a*).

### Confocal microscopy

A Zeiss Laser Scanning Microscope-780 or 880 (Zeiss) was used to generate the fluorescent images. Settings were optimized to visualize GFP, YFP (laser wavelength, 488 nm; detection wavelength, 493 to 598 nm) or chlorophyll (laser wavelength, 545 nm; detection wavelength, 604 to 694 nm). All direct comparisons of images were performed with the same settings. Maximum intensity projections were generated from z-stacks of multiple tiles to visualize the entire apical region of the gynoecium/silique. Samples were mounted on a glass slide with 1% low-melt agarose (Bio-Rad) and submerged in de-ionized water.

### Use of publicly available genomics data

M-values for the expression of all genes detected during flower development were previously generated (Ryan *et al*., 2015). Transcripts per million for each biological replicate and time-point from Kivivirta *et al*., 2020 were generously shared with us by Dr. Annette Becker and Clemens Rössner (Justus-Liebig Universität Giessen, Germany). ChIP-Seq peaks were visualized using online software at ChIP-Hub (https://biobigdata.nju.edu.cn/ChIPHub/) (Fu *et al*., 2022). Seq-DAP-seq sequences were downloaded from the genome browser (https://genome.ucsc.edu/s/ArnaudStigliani/MADS) (Lai *et al*., 2020).

### Scanning electron microscopy

Samples were fixed and processed for SEM as previously described in Laux *et al*., 1996. Briefly, samples were harvested in 1 x PBS (P4417-100TAB, Sigma-Aldrich) at room temperature. The PBS was then replaced with a 4% solution of 50% glutaraldehyde (4995.1, Roth) in 1 x PBS and incubated overnight at 4°C. A dehydration series was then performed using freshly prepared solutions of 30%, 50%, 70%, and 90% ethanol (v/v) at room temperature for 30 min each while being gently rocked. Samples were then incubated in 100% ethanol for 60 min at room temperature. Ethanol was then replaced with 100% ethanol and placed at 4°C. A Leica CPD300 or Quorum K850 were used for critical point drying and samples were coated with gold using an Emitech SC 7640 (Polaron) or Quorum Q150T sputter coater. Samples were imaged using a Hitachi TM 4000Plus or a Supra 40VP scanning electron microscope (Zeiss).

### Chemical treatments

A solution containing 10 µM dexamethasone (DEX; Sigma) and 0.015% Silwet L-77 in deionized water was applied liberally to the inflorescences of plants with a Pasteur pipette. Mock treatments contained all the same components without dexamethasone. Ethanol vapor treatments were performed by sealing the plants in containers (18 cm × 32 cm × 50 cm) with clear plastic lids and two 50 mL tubes that each contained 10 mL of 100% ethanol for 6 h.

### Reverse transcription quantitative PCR

Samples were harvested on liquid nitrogen and stored at −80°C until processed. Harvesting of gynoecia between stages 10-13 was done under a stereomicroscope from the primary inflorescences of 15-20 plants. Only gynoecia, where the majority of the stigmatic tissue was clear of pollen (i.e. largely unfertilized), were harvested. Harvesting of stage 13 gynoecia from the *AG-amiRNA*^*i*^ line for the time-course (Figure 3C) was done by initially removing all flowers and siliques that were beyond stage 13. Then flowers at anthesis were harvested for the 0 d time-point. Plants were treated with DEX and 1 d later, 2-3 flowers from the same plants at anthesis were harvested on liquid nitrogen. This continued until 7 d after the treatment from 15-20 plants.

**Figure 2.**
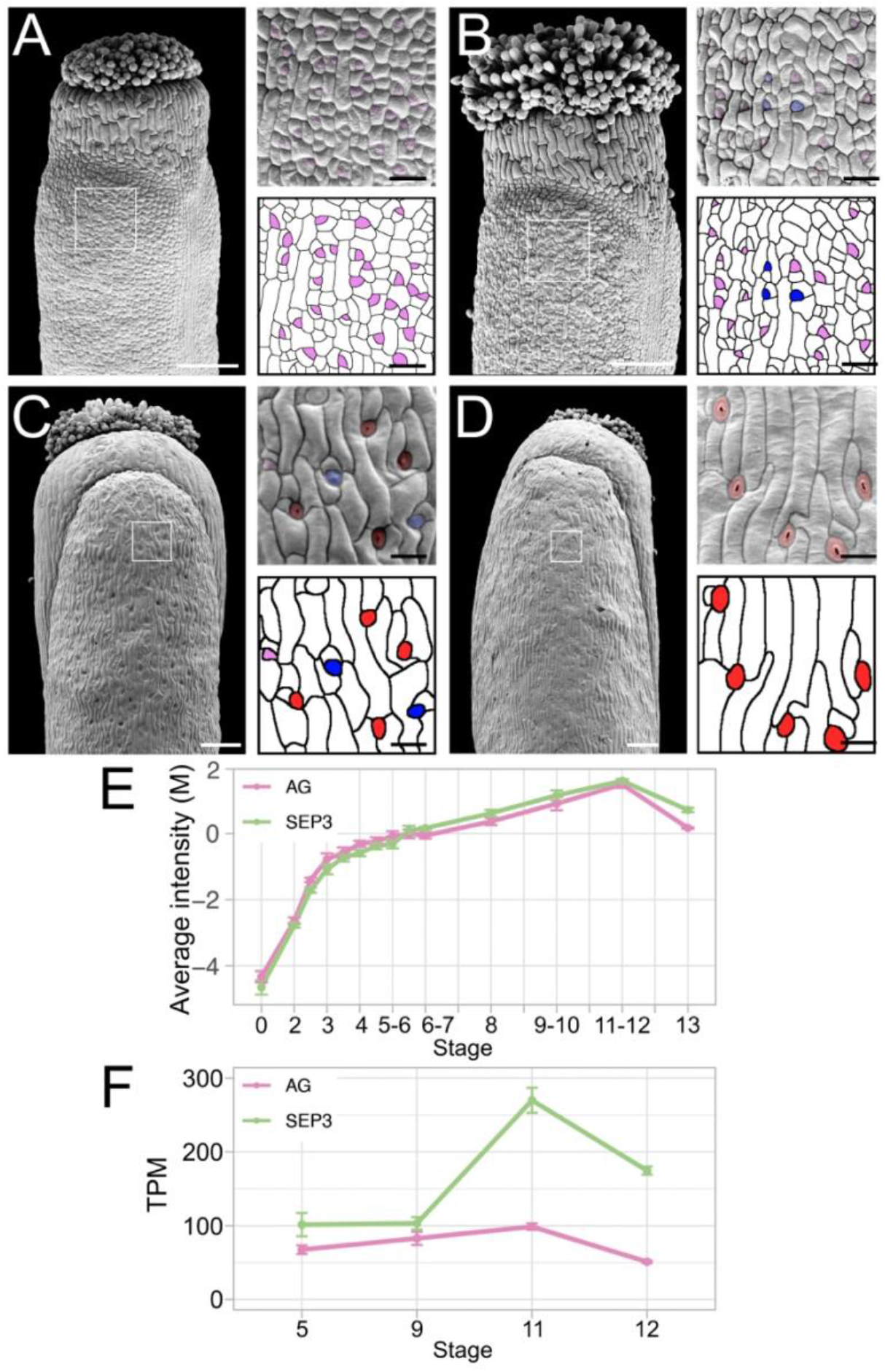
Progression of stomatal development on gynoecial and silique valves. (A-D) Scanning electron micrographs of gynoecia at floral stage (A) 12, (B) 13, (C) 15-16 gynoecia and (D) a stage >17 silique with magnifications of the cell surface (from white box). Scale bars for images of whole gynoecia are 100 µm. Scale bars for magnifications are 20 µm. Purple, blue, and red highlights indicate early, mid, and late stage stomatal lineage morphology, respectively. (E-F) Levels of *AG* and *SEP3* mRNAs over the course of (E) flower development (Ryan *et al*. 2015) and (F) gynoecium development (Kivivirta *et al*. 2020). Error bars in (E-F) are s.e.m of three and four biological replicates, respectively. M, log-ratio of time-point to a reference; TPM, transcripts per million.

**Figure 3.**
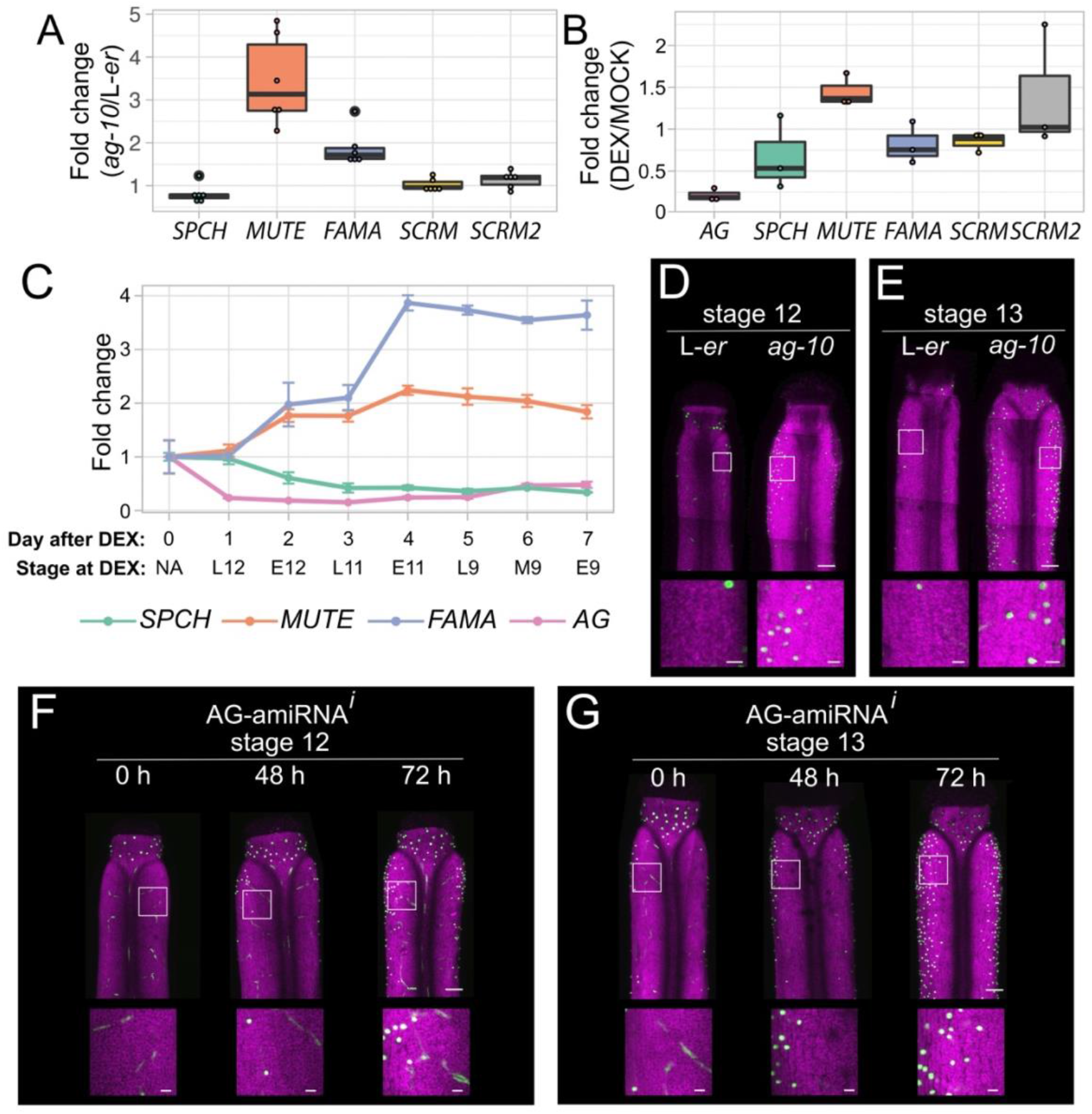
Transcriptional response of master regulators of stomatal development to perturbation of *AG*. (A-B) Levels of mRNAs encoding stomatal bHLH transcription factor regulators, as determined by RT-qPCR, in (A) *ag-10* stage 10-13 gynoecia relative to L-*er* stage 10-13 gynoecia, (B) dexamethasone-treated *AG-amiRNA*^*i*^ (*OPpro:AG-amiRNA/35Spro:GR-LhG4*) in stage 10-13 gynoecia relative to mock-treated *AG-amiRNA*^*i*^ in stage 10-13 gynoecia 24 h after treatments. Each dot in (A-B) represents the mean of an individual biological replicate. (C) Levels of *SPCH, MUTE, FAMA*, and *AG* mRNAs, as determined by RT-qPCR, in stage 13 gynoecia after treatment with dexamethasone relative to untreated (0 d) in stage 13 gynoecia. “Day after DEX” indicates the number of days that gynoecia were treated with DEX before being harvested at anthesis (stage 13), with “0 d” representing the untreated sample. “Stage at DEX” indicates the approximate stage of the flower/gynoecium when DEX treatment was applied. Error bars are s.e.m of three biological replicates. (D-G) Maximum intensity projections of stitched confocal laser scanning z-stack micrographs of (D, F) stage 12 and (E, G) stage 13 gynoecia from plants harbouring a *FAMApro:2xYFP* transgene in (D-E) L-*er* and *ag-10* backgrounds, and (F-G) the *AG-amiRNA*^*i*^ (*OPpro:AG-amiRNA/35Spro:GR-LhG4*) background before treatment (0 h) and after dexamethasone treatment (48 h and 72 h). YFP is coloured green and chlorophyll fluorescence is coloured magenta. Scale bars for images of whole gynoecia are 100 µm. Scale bars for insets are 20 µm.

Samples were disrupted using a TissueLyser II (Qiagen) with adapters that were kept at −80°C. Total RNA was extracted from the samples using an RNeasy Plant Mini kit (Qiagen) or a Spectrum Plant Total RNA kit (Sigma-Aldrich). Removal of genomic DNA was performed with the Turbo DNA-free Kit (Invitrogen) or the On-Column DNase I Digestion set (Sigma-Aldrich). Reverse transcription was performed with the SuperScript™ IV First-Strand Synthesis System (Invitrogen) or RevertAid H Minus Reverse Transcriptase (ThermoFisher Scientific). Quantitative PCR was performed with either iQ SYBR green (Bio-Rad) or PowerUp SYBR Green (Thermofisher Scientific) using a Lightcycler 480 (Roche) or Stratagene Mx3005 (Agilent Technolgies). At least 2 technical measurements for each biological replicate were made and the mean of these technical replicates was used to represent each biological replicate. The number of biological replicates for each RT-qPCR experiment is indicated in Supplemental Table 3. Melting curves were obtained for the reactions, revealing single peak melting curves for the amplified products. The amplification data were analyzed using the second derivative maximum method, and resulting values were converted into relative expression values using the comparative cycle threshold method (Livak and Schmittgen, 2001). Primers used in this study are listed in Supplemental Table 2.

### Motif enrichment analysis

The 1421 sequences identified as bound by AG in Ó’Maoiléidigh *et al*., 2013 were downloaded from www.araport.org (Pasha *et al*., 2020). These sequences were uploaded to the MEME-ChIP interface on meme-suite.org (Machanick and Bailey, 2011; Bailey *et al*., 2015). The settings applied and motifs identified are summarized in Supplemental Table 4.

### Genomic sequence alignents

The genomic and inferred amino acid sequences of *MUTE* were downloaded from The Arabidopsis Information Resource (www.arabidopsis.org) (Berardini *et al*., 2015). Orthologs of *AtMUTE* were previously identified in *Arabidopsis lyrata, Capsella rubella*, and *Eutrema salsugineum* (formerly *Thellungiella halophila*) (Ran *et al*., 2013). We identified putative orthologs in *Capsella grandiflora, Arabidopsis halleri*, and *Boechera stricta* by using the TBLASTN functionality in Phytozome (https://phytozome-next.jgi.doe.gov/) with the AtMUTE amino acid sequence against the above genomes (Goodstein *et al*., 2012). We found one clear candidate in each genome, which included the same orthologs identified in Ran *et al*., 2013. The TBLASTN search results are presented in Supplemental Table 5. Sequence alignments were performed with mVISTA (https://genome.lbl.gov/vista/mvista/submit.shtml) using standard settings (Frazer *et al*., 2004).

### Statistical analysis

Some recommendations from Wasserstein *et al*., 2019 are implemented here. The term “statistically significant” has been avoided and *p*-values have been reported as continuous where possible. Although grouping data based on statistical thresholds is not recommended (Wasserstein *et al*., 2019), a grouping threshold is provided in supplemental tables as a helpful guide, but results are not interpreted on the basis of these groupings. Data were analyzed using the R studio software (Team, 2022). For multiple comparisons, data were analyzed using a one-way analysis of variance (ANOVA). Post hoc tests were employed if a significant result was obtained from the above tests (p < 0.01). The base function pairwise.t.test was used following an ANOVA test, with analysis being paired or unpaired depending on the experimental design. Adjustments for multiple corrections were performed in each case using the Benjamini–Hochberg method (Benjamini and Hochberg, 1995). For the box plots, the box encapsulates the 25th to 75th percentile, and the error bars encapsulate the 10th and 90th percentiles. The horizontal line running through the box indicates the median, and each point represents an individual plant.

### Gel shift assays

Electrophoretic mobility shift assays were performed essentially as described by Melzer *et al*., 2009. DNA probes were ordered as single stranded oligo, annealed, and radioactively labelled with [α-P^32^] dATP by a Klenow fill-in reaction of 5′-overhangs. Probe sequences are shown in Supplementary Table 6. Proteins were produced *in vitro* via the TNT SP6 Quick Coupled Transcription/Translation System (Promega) following manufacturer’s instructions. Plasmids for *in vitro* transcription/translation of SEP3 and SEP3ΔC have been generated previously (Melzer, et al. 2009). For generation of the plasmids for *in vitro* transcription/translation of AG, SHP1, and SHP2, coding sequences of *AG* (X53579.1), *SHP1 (NM_115740*.*3)*, and *SHP2* (NM_129844.5) were synthesized via the GeneArt gene synthesis service (Thermo Fisher Scientific) and cloned into pTNT (Promega) using *Eco*RI and *Sal*I restriction sites. The composition of the protein-DNA binding reaction buffer was essentially as described by Egea-Cortines *et al*., 1999 with final concentrations of 1.6 mM EDTA, 10.3 mM HEPES, 1mM DTT, 1.3 mM spermidine hydrochloride, 33.3 ng µl^−1^ Poly dI/dC, 2.5 % CHAPS, 4.3 % glycerol, and 6 µg µl^−1^ BSA. For each binding reaction 2 µl of *in vitro* transcribed/translated protein were co-incubated with 0.1 ng of radioactively labelled DNA probe in a total volume 12 μl. Binding reactions were incubated overnight at 4°C and loaded on a polyacrylamide (5% acrylamide, 0.1725% bisacrylamid) 0.5× TBE gel. The gel was run at room temperature with 0.5× TBE buffer for 3 h at 7.5 V cm^−1^, and afterwards vacuum-dried and exposed onto a phosphorimaging screen to quantify signal intensities. For size comparison a radioactively labelled DNA ladder (100 bp DNA ladder, New England Biolabs) was applied.

## Results

### Stomatal development on gynoecial valves

As mentioned above, the photosynthetic activity of siliques of the Brassicaceae family contributes significantly to the carbon requirements of seeds (Hua *et al*., 2012; Wang *et al*., 2016; Zhu *et al*., 2018). Siliques assimilate atmospheric CO_2_ (Sheoran *et al*., 1991; Gammelvind *et al*., 1996; Hua *et al*., 2012), probably through the stomata present on the valve epidermis, which are not present on pre-anthesis gynoecial valves (Supplemental Figure 1B) (Smyth *et al*., 1990). The presence of stomata on floral organs was previously reported (Smyth *et al*., 1990; Geisler *et al*., 1998; Nadeau and Sack, 2002*a*) (Supplemental Figure 1C-E), however, the progression of stomatal development on these organs has not been fully described. We addressed this knowledge gap by examining the developmental progression of stomata on the siliques of *A. thaliana*.

We started by surveying the mRNA levels of master regulators of stomatal development as reported by publicly available transcriptomics datasets of flower and gynoecium development (Figure 1A-B). We focused on the expression of three bHLH transcription factor-coding genes, *SPCH, MUTE*, and *FAMA*, which are necessary and sufficient to promote different stages of the stomatal lineage (Ohashi-Ito and Bergmann, 2006; MacAlister *et al*., 2007; Pillitteri *et al*., 2007). A distinct peak of *SPCH* and *MUTE* mRNA was detected at stage 6-7 and 8 of flower development, respectively, in genome-wide expression profiling using a synchronous flowering system (Figure 1A) (Ryan *et al*., 2015). These peaks probably corresponded to the initiation and progression of the stomatal lineage on sepals, which began to mature at approximately stage 10 (Supplemental Figure 1C). Levels of both *SPCH* and *MUTE* mRNA then decreased until anthesis, although *MUTE* levels plateaued between stages 11 and 13. In contrast, *FAMA* levels continued to increase as flower development progressed (Figure 1A). A more recent transcriptomics datasets derived from laser-microdissected gynoecia at different stages of flower development provided improved spatial resolution (Kivivirta *et al*., 2020). These data are not complicated by the presence of other floral organs that bear stomata, such as the sepals and stamens, although the latest stage analyzed was stage 12 of flower development. *SPCH* mRNA levels were relatively high from approximately stage 5 and rose slightly until stage 11 and then decreased (Figure 1B). *MUTE* mRNA was not detectable before stage 11 when it was weakly expressed and then increased ~3-fold by stage 12 (Figure 1B). *FAMA* mRNA was also not detected at stage 5 but increased steadily by ~8-fold between stages 9 and 12 (Figure 1B). Based on these data, we concluded that *MUTE* and *FAMA* transcription in the gynoecium was initiated at approximately stage 11-12, whereas *SPCH* transcription was initiated earlier.

To cross-examine these transcriptomic data and to provide a better resolution of the expression of these genes in the gynoecium, we obtained transgenic lines that harbor fluorescent translational or transcriptional reporters for *SPCH* (Davies and Bergmann, 2014), *MUTE* (Pillitteri *et al*., 2007), and *FAMA* (Adrian *et al*., 2015) whose transcription is driven by their endogenous regulatory elements (Figure 1C-E). Using laser-scanning confocal microscopy, we could not detect SPCH-YFP in stage 11 gynoecial valves, but the SPCH-YFP protein was abundant in the epidermis of late stage 12 and stage 13 gynoecial valves (Figure 1C). The contrast between mRNA and protein accumulation may indicate the presence of a post-translational mechanism to control SPCH protein levels (Figure 1B, C). A pattern of protein accumulation was found for MUTE-GFP, which was largely in agreement with the transcriptomics data in gynoecia (Figures 1B and 1D). The accumulation of YFP from the *FAMApro:NLS-2xYFP* transgene was delayed relative to either of these reporters, with few fluorescent foci present at stage 12, which became slightly more abundant at stage 13 (Figure 1E). At stage 14, accumulation of YFP was observed as fluorescent foci throughout the gynoecial valve (Figure 1E). Therefore, the SPCH and MUTE proteins accumulate after stage 11 but before late stage 12, whereas *FAMA* transcription is initiated after late stage 12.

We then examined the formation of stomatal cell lineage types on gynoecial valves using scanning electron microscopy (SEM). We categorized the stomatal lineage into ‘early’ (including meristemoids and young GMCs), ‘mid’ (including GMCs before signs of asymmetric cell division were observed), and ‘late’ stages (including GMCs where asymmetric cell division had started through to mature stomata). We observed early-stage stomatal lineage cells on gynoecial valves during stage 12 of flower development whereas more mature stomatal cell lineages were largely absent (Figure 2A, Supplemental Table 7). At stage 13, early-stage lineage cells remained most common, however, mid-stage cells became more prevalent and late-stage cells were occasionally observed (Figure 2B, Supplemental Table 7). Post-fertilization, at stage 15-16, the frequency of early-stage cells reduced dramatically with a commensurate increase in mid-stage and late-stage cells (Figure 2C, Supplemental Table 7), until most stomatal lineage cells were of late-stage on older (>stage 17) siliques (Figure 2D, Supplemental Table 7). Stomata also formed on the style of the gynoecium prior to the formation of stomata on the valves, with mature stomata present on the style of stage 12 gynoecia (Supplemental Figure 1E). However, the style comprises only a small fraction of the total area of the mature silique (Supplemental Figure 1F), suggesting that stomata on the style contribute only a small fraction of the total carbon fixation and transpiration capacity of the silique.

Given the close association between fertilization and progression of stomatal development, we tested whether a dependent relationship existed (Supplemental Figure 2). To this end, we emasculated flowers before anthesis and examined gynoecial valves 5 d after anthesis by SEM (Supplemental Figure 2A). We found mature stomatal complexes on these unfertilized gynoecial valves (Supplemental Figure 2B), which indicated that maturation of stomatal complexes on the gynoecium does not depend on fertilization.

### Expression of *AG* and *SEP3* during the formation of stomatal complexes

The C class protein AG controls the specification and development of stamens and carpels by forming dimers or tetramers with the E class protein SEP3 (Bowman *et al*., 1989; Pelaz *et al*., 2000*a*; Honma and Goto, 2001; Hugouvieux *et al*., 2018). Notably, it has been shown that AG suppresses the formation of trichomes on carpel valves by controlling the expression of other developmental regulators (Ó’Maoiléidigh *et al*., 2013, 2018). In addition, trichomes form on the gynoecia of plants with reduced *SEP* activity (Ditta *et al*., 2004). Because stomata, like trichomes, are formed by the epidermis, we hypothesized that an AG-SEP3 complex might also repress the formation of stomata on carpel valves. To explore this, we first assessed published expression data for *AG* and *SEP3* in gynoecial tissues over the course of stomatal initiation and maturation (Figure 2E-F, Supplemental Figure 3). *In situ* hybridizations showed *AG* mRNA accumulation throughout the carpels during stage 8 before its expression becomes restricted and absent from stage 12 gynoecial valves (Bowman *et al*., 1991*a*). A similar expression pattern was observed of *SEP3* mRNA in the gynoecium, although its expression was restricted from gynoecial valves from approximately stage 10 (Mandel and Yanofsky, 1998). This is consistent with the results of transcriptomics experiments, where *AG* and *SEP3* mRNA levels begin to decrease from approximately stage 11 of flower development (Figure 2E-F) (Ryan *et al*., 2015; Kivivirta *et al*., 2020).

AG-GFP and SEP3-GFP protein accumulation was analyzed in transgenic plants that harbored *AGpro:AG-GFP* and *SEP3pro:SEP3-GFP* transgenes that drive transcription from the corresponding endogenous regulatory elements (Urbanus *et al*., 2009). AG-GFP fluorescence was observed throughout stage 12 gynoecial valves in these plants even though *AG* mRNA is not present at this stage (Bowman *et al*., 1991*a*), possibly due to the low turnover of AG at these stages (Urbanus *et al*., 2009). We verified these results for AG-GFP using a similar but independently generated *AGpro:AG-GFP* transgenic line using laser-scanning confocal microscopy (Supplemental Figure 3A-B) (Ó’Maoiléidigh *et al*., 2013). We also imaged stage 13 gynoecia and found that fluorescent signal stemming from AG-GFP was more intense in the replum and style relative to the valves (Supplemental Figure 3C). Additionally, SEP3-GFP protein accumulated to much higher levels in the replum and valve margins at stage 12 when compared to the gynoecial valves (Urbanus *et al*., 2009). Taken together with the transcriptomics data, it appears that the abundance of AG and SEP3 proteins in the gynoecial valves reduces significantly between stages 11 and 13 of flower development.

### Transcriptional response of master regulators of stomatal development to *AG* perturbation

To test whether a perturbation of *AG* activity would result in the differential expression of master regulators of stomatal development, we harvested stage ~10-13 gynoecia from L-*er* wild-type plants harboring a fully functional copy of *AG* and those that were homozygous for the weak *ag-10* mutation (Liu *et al*., 2011). Plants that are homozygous for *ag-10* form gynoecia that are relatively normal in appearance, although they tend to bulge (Liu *et al*., 2011). We extracted total RNA from these plants and using reverse-transcription combined with quantitative PCR (RT-qPCR), we found that the mean mRNA levels of *MUTE* and *FAMA* were increased in *ag-10* carpels to ~3.4 (*p* = 0.0002, two-tailed paired t-test) and ~1.9-fold (*p* = 0.0004, two-tailed paired t-test), respectively, relative to wild-type counterparts (Figure 3A, Supplemental Table 3). In contrast, the mean levels of *SPCH* mRNA were mildly decreased to 0.8-fold (*p* = 0.09, two-tailed paired t-test) while the mean mRNA levels of *SCRM* and *SCRM2* appeared unchanged (Figure 3A, Supplemental Table 3). Next, we used a transgenic line that harbors a dexamethasone (DEX) inducible artificial microRNA that is designed to target the *AG* mRNA (*AG-amiRNA*^*i*^) (Ó’Maoiléidigh *et al*., 2013). We treated the inflorescences of these plants with a DEX-containing solution and a mock solution, and harvested stage ~10-13 gynoecia 24 h later. By RT-qPCR, we found that mean *MUTE* mRNA levels were increased to ~1.4-fold (*p* = 0.08, two-tailed paired t-test) in DEX-treated samples relative to the mock control (Figure 3B, Supplemental Table 3), consistent with the increased expression detected in *ag-10*. None of the other bHLH-coding genes were consistently differentially expressed (Figure 3B, Supplemental Table 3).

We then used the *AG-amiRNA*^*i*^ line in a time-course experiment where we harvested stage 13 gynoecia of untreated plants (0 d) and then treated the inflorescences of plants with a DEX-containing solution. We harvested only gynoecia that had reached anthesis every day for seven days following that treatment. Therefore, gynoecia that were harvested 1 d following DEX-treatment would have been ~late-stage 12 gynoecia at the time of DEX treatment. Gynoecia harvested at 2 d following DEX-treatment would have been ~early stage 12 gynoecia, and so on (Figure 3C). Following harvesting, we processed the samples and determined mRNA levels of *SPCH, MUTE, FAMA*, and *AG* using RT-qPCR. One day after treatment, there was very little change in the levels of *SPCH, FAMA*, or *MUTE* mRNAs whereas *AG* mRNA was reduced to 0.2-fold when compared with 0 d (*p*.*adj* < 10^−9^, pairwise two-tailed t-tests), as has been previously described (Figure 3C, Supplemental Table 3) (Ó’Maoiléidigh *et al*., 2013). The lack of response from the bHLH-encoding genes may have been because the *AG-amiRNA* was activated in late-stage 12 gynoecia when *AG* mRNA levels were already very reduced in the gynoecial valve, although *AG* mRNA continued to accumulate in other tissues (Bowman *et al*., 1991*a*). At 2 d post-induction, which roughly corresponded to activation of the *AG-amiRNA* in early stage 12 gynoecia, the mRNA levls of both *MUTE* and *FAMA* increased to ~1.8 (*p*.*adj* = 0.006, pairwise two-tailed t-tests) and ~2-fold (*p*.*adj* = 0.02, pairwise two-tailed t-tests) relative to 0 d, respectively, as *SPCH* mRNA levels decreased to 0.6-fold (*p*.*adj* = 0.006, pairwise two-tailed t-tests) (Figure 3C, Supplemental Table 3). *MUTE* and *FAMA* expression then continued to increase until 4 d post-induction where they peaked at ~2.2 (*p*.*adj* = 0.0004, pairwise two-tailed t-tests) and ~3.9-fold (*p*.*adj* < 10^−5^, pairwise two-tailed t-tests) relative to 0 d, respectively, and then began to decrease. At 5 d post-induction, *AG* mRNA levels began to recover, which correlated with a reduction in *MUTE* mRNA levels, which *FAMA* partially mimics, until the end of the experiment at 7 d post induction (Figure 3C, Supplemental Table 3). Taken together, we concluded that AG represses the expression of *MUTE*, and *FAMA* during gynoecium development.

To improve our understanding of the spatial resolution with which AG controls the expression of these stomatal genes, we introduced the *FAMApro:2xYFP* reporter into the *ag-10* background and imaged gynoecia at late stage 12 and stage 13 using confocal laser scanning microscopy (Lee *et al*., 2019). We observed very few fluorescent foci on the valves of stage 12 wild-type gynoecia and stage 13 valves (Figure 3D-E, Supplemental Table 8), as previously described for the *FAMApro:NLS-2xYFP* reporter (Figure 1E). In contrast, over 20 times the number of foci were observed on the valves of stage 12 (*p* = 0.06, two-tailed paired t-test) and stage 13 *ag-10* gynoecia (*p* = 0.01, two-tailed paired t-test) (Figure 3D-E). We then introduced the *FAMApro:2xYFP* reporter into the *AG-amiRNA*^*i*^ transgenic line. We treated inflorescences with a DEX-containing or mock solution and imaged stage 12 and 13 gynoecia every day for 3 days (Figure 3F-G, Supplemental Figure 4, Supplemental Table 8). Similar to the *ag-10* observations (Figure 3D-E) and mirroring the results of the RT-qPCR experiments (Figure 3C), fluorescent foci were prevalent on valves of the DEX-treated gynoecia at 48 and 72 h (Figure 3F-G). Far fewer fluorescent foci were observed on the valves of untreated samples (0 h, Figure 3F-G), mock-treated counterparts (Supplemental Figure 4B, D), or DEX-treated samples after 24 h (Supplemental Figure 4A, C). Statistical summaries of these *AG-amiRNA*^*i*^ data are presented in Supplemental Table 8. These confocal data support the observations of the RT-qPCR experiments and clarify that expression of *FAMA* is not elevated throughout the gynoecial epidermis but only in a pattern that is consistent with stomatal formation on the gynoecial valves.

**Figure 4.**
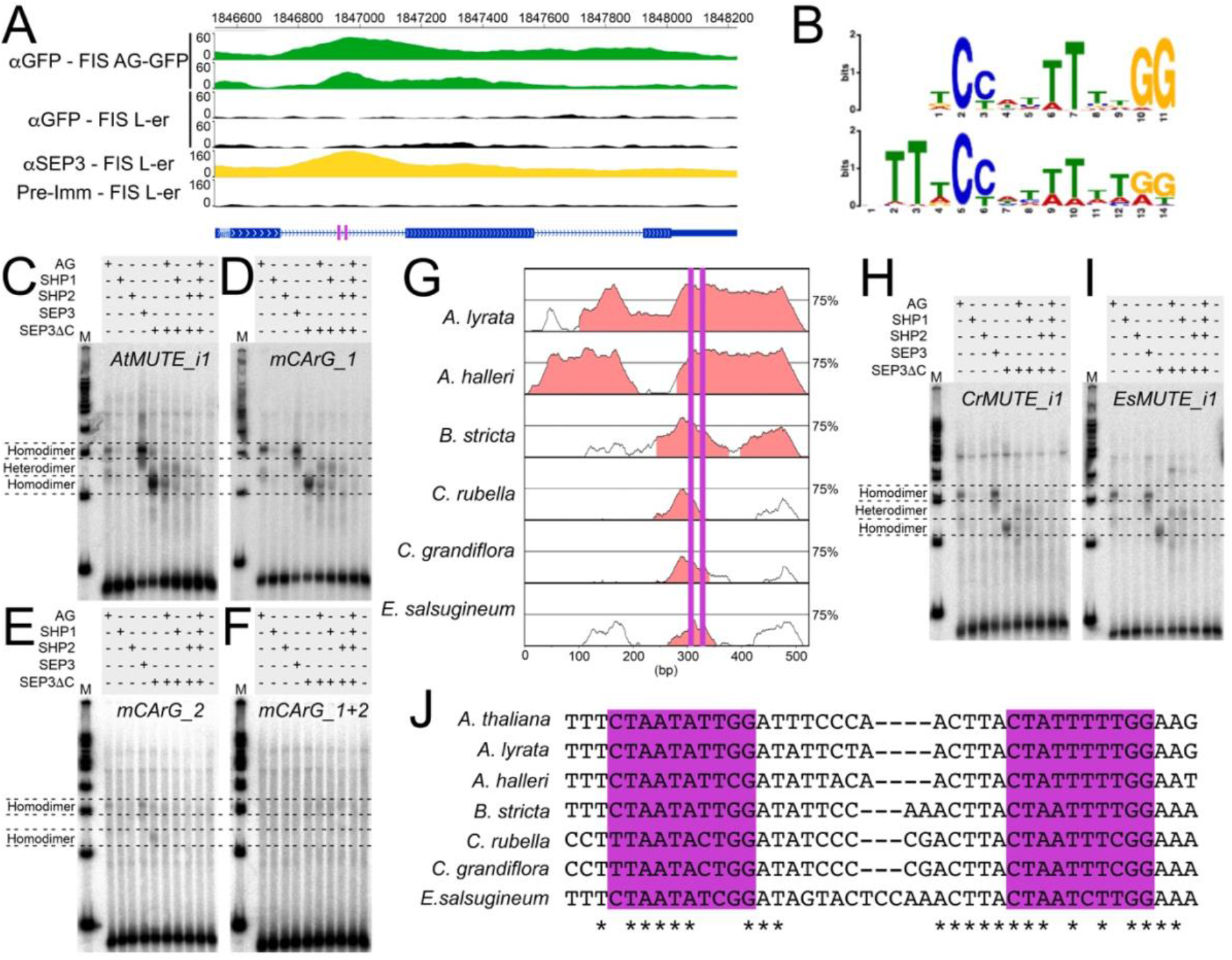
Interaction between AG, SHP1, SHP2, SEP3, and first intron of *MUTE*. (A) Tracks indicating enrichment of sequences in the first intron of *MUTE* from two replicates of ChIP-Seq of AG-GFP (green tracks) and SEP3 (yellow tracks, combined results of two replicates), proteins in the floral induction system (FIS) (Ó’Maoiléidigh *et al*., 2013; Pajoro *et al*., 2014). Corresponding control ChIP-Seq experiments are coloured in black. A schematic of the gene structure of *MUTE* blue is below (exon, rectangles; introns, lines). Two CArG motifs were identified within the first intron of *MUTE* (purple rectangles). (B) Binding logos from MEME and STREME analyses of 1421 binding sites identified in a ChIP-Seq of AG (Ó’Maoiléidigh *et al*., 2013). (C-F) Protein-DNA gel shift assays using combinations of AG, SEP3, SEP3ΔC, SHP1, and SHP2 recombinant protein and (C) a wild-type *AtMUTE* probe (*AtMUTE_i1*), (D) a probe where CArG_1 is mutated (*mCArG_1*), (E) a probe where CArG_2 is mutated (*mCArG_2*), and (F) a probe where both CArG motifs are mutated (*mCArG_1+2*). M, molecular weight marker. (G) An mVISTA alignment of the first intron of *MUTE* from various members of the Brassicaceae relative to *A. thaliana*. Regions highlighted in salmon have been designated as “Conserved Non-Coding Sequences” by mVISTA. Purple lines indicate the position of each CArG motif. (H-I) Protein-DNA gel shift assays using combinations of AG, SEP3, SEP3ΔC, SHP1, and SHP2 protein and (H) a wild-type *CrMUTE* probe (*CrMUTE_i1*) and (I) a wild-type *EsMUTE* probe (*EsMUTE_i1*). M, molecular weight marker. (J) Sequence alignments using some of the conserved region identified in (B) containing both CArG motifs. Asterisks indicate conserved nucleotide and the two CArG motifs are highlighted by purple rectangles. Unadjusted images for both replicates of the protein-DNA gel shift assays can be found in Supplemental Figure 8.

### Interaction between AG, SHP, SEP3, and the first intron of *MUTE*

*AG* activity is negatively correlated with stomatal development on gynoecial valves and perturbation of *AG* results in elevated mRNA levels of key stomatal regulators, such as *MUTE* and *FAMA*. To understand whether the regulation of *MUTE* and *FAMA* transcription was directly controlled by AG, we interrogated chromatin immunoprecipitation followed by next-generation sequencing (ChIP-Seq) data available for AG (Ó’Maoiléidigh *et al*., 2013). We found that AG bound to the first intron of *MUTE* (Figure 4A), which was previously identified as a direct target (Ó’Maoiléidigh *et al*., 2013). *MUTE* was also identified as a putative direct target of SEP3 (Pajoro *et al*., 2014) (Figure 4A), which forms heterodimers and quaternary complexes with AG to control floral organ development (Honma and Goto, 2001; Theißen *et al*., 2016; Hugouvieux *et al*., 2018). Within the first intron of *MUTE* are two motifs with similarity to CArG motifs, to which these MADS domain transcription factors bind (Wuest *et al*., 2012; Ó’Maoiléidigh *et al*., 2013; Pajoro *et al*., 2014). Analysis of the genomic sequences bound by AG revealed an enrichment of CArG motifs, which were very similar to the motifs identified in the first intron of *MUTE* (Figure 4B, Supplemental Table 4). *MUTE* was also identified as a direct target of the AG-SEP3 complex through sequential immunoprecipitation followed by DNA affinity purification and sequencing experiments (seq-DAP-Seq) (Supplemental Figure 5) (Lai *et al*., 2020).

**Figure 5.**
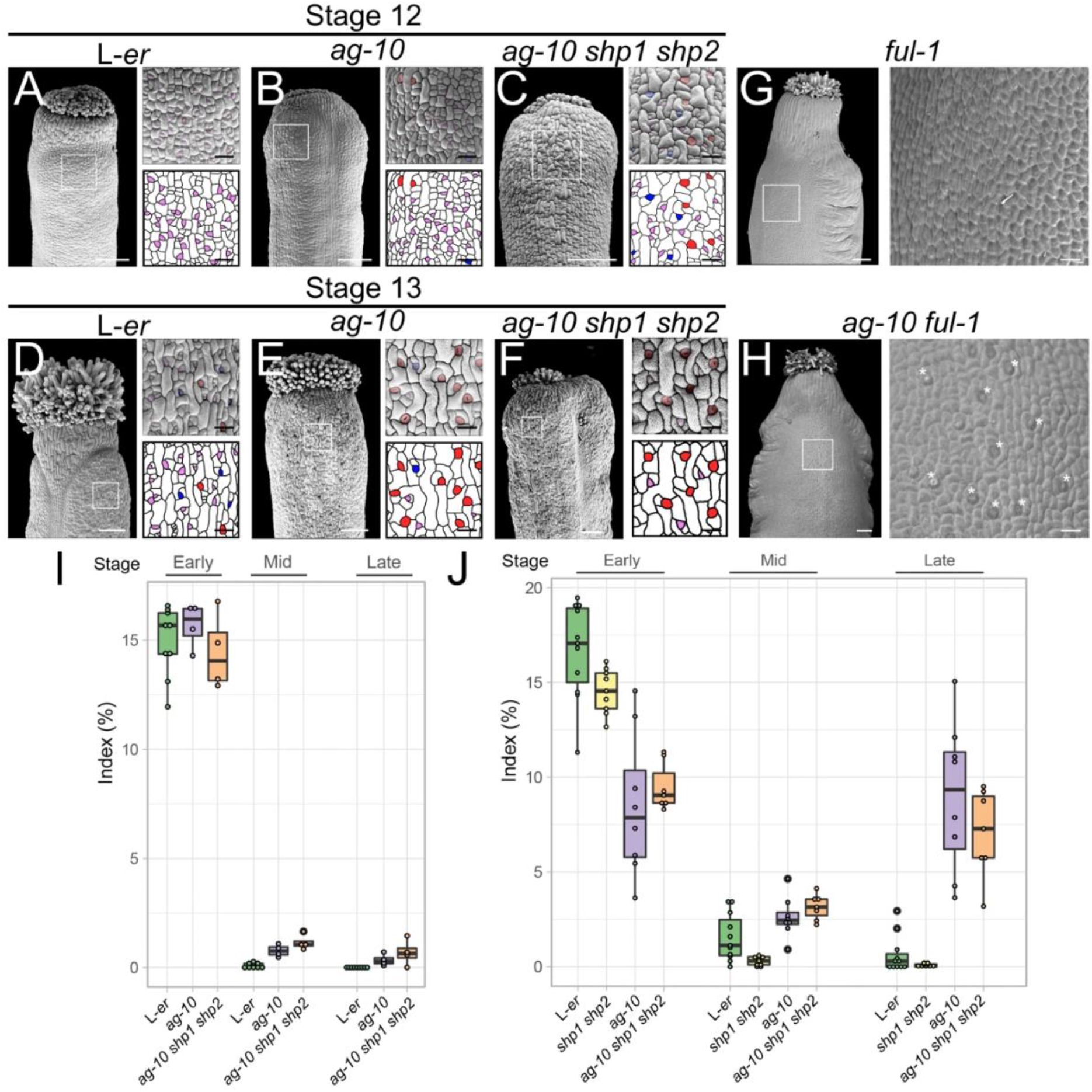
Stomatal development on gynoecial valves in response to *AG* and *SHP* perturbation. (A-F) Scanning electron micrographs of (A-C) stage 12 and (D-F) stage 13 L-*er, ag-10*, and *ag-10 shp1 shp2* gynoecia, as indicated. Purple, blue, and red highlights indicate early, mid, and late-stage stomatal lineage morphology, respectively. (G-H) Scanning electron micrographs of a (G) *ful-1* and (H) *ag-10 ful-1* silique. Asterisks indicate presence of stomata. Scale bars for images of whole gynoecia are 100 µm. Scale bars for magnifications are 20 µm. (I-J) Index of early, mid, and late stomatal lineages based on morphology from scanning electron micrographs from (I) stage 12 and (J) stage 13 L-er, *shp1 shp2, ag-10, ag-10 shp1 shp2* gynoecia. Each dot represents an individual sample.

We performed gel shift assays to cross-validate these experiments and to determine whether both CArG motifs in the *MUTE* first intron are required for AG and SEP3 binding. We synthesized probes with the same sequence as a region of the *MUTE* first intron that contains the two CArG motifs (*AtMUTE_i1*) and incubated them with the AG and SEP3 proteins (Supplemental Table 6, Figure 4C). We also used a truncated version of SEP3 (SEP3ΔC), which has been shown to retain activity (Melzer *et al*., 2009), to visualize shifts between homodimers of SEP3 and AG-SEP3 heterodimers. We found that incubation of SEP3, SEP3ΔC, or AG individually resulted in the appearance of a single retarded fraction of the DNA probe, suggesting that homodimers of each of these proteins can bind the *MUTE* first intron (Figure 4C). In accordance to the similar size of the full-length proteins, the retarded fractions produced by SEP3 and AG migrated with approximately the same speed, whereas the retarded fraction produced by SEP3ΔC migrated faster, likely due to the reduced protein size. Co-incubation of AG with SEP3ΔC resulted in the appearance of an additional retarded fraction of intermediate electrophoretic mobility, probably corresponding to AG-SEP3ΔC heterodimers. We also incubated the SHP1 and SHP2 paralogs of AG with SEP3ΔC in separate reactions and observed a shifted band when SHP1 was incubated with SEP3ΔC at a height similar to that of the putative AG-SEP3ΔC heterodimers. Only a very weak band was present to suggest that a SHP2-SEP3ΔC heterodimer could bind the *AtMUTE_i1* probe and there was very weak support for SHP1 or SHP2 homodimer binding. Furthermore, incubation of all 4 proteins together did not result in a shift, suggesting that a multimeric complex does not form at the *MUTE* intron (Figure 4C).

Next, we tested whether the CArG motifs present in this region were required for binding. We incubated combinations of the above proteins with probes harboring modifications of the CArG motifs individually (*mCArG_1, mCArG_2*) or simultaneously (*mCArG_1*+*2*), which were intended to disrupt protein binding (Supplemental Table 6). Similar results to the *AtMUTE_i1* probe were obtained when *mCArG_1* was incubated with the same combinations of proteins as above (Figure 4D). However, incubation with the *mCArG_2* probe abolished the formation of bands corresponding to heterodimer formation. Furthermore, the formation of bands corresponding to homodimers were reduced significantly (Figure 4E). Disruption of both CArG motifs (*mCArG_1+2*) eliminated all shifted bands (Figure 4F). This suggests that CArG_2 interacts with heterodimers and that both CArG motifs can interact with homodimers, although the tested proteins have the highest affinity for CArG_2.

We compared the sequence of the first intron of *A. thaliana MUTE* to the sequences of the first intron from orthologs of *MUTE* from other members of the Brassicaceae family (Supplemental Table 5). We found that a region of approximately 60 bp was well-conserved, which contained both CArG motifs (Figure 4G). For CArG_2, the AT track in the center of the CArG motif was well maintained in four species. However, a single T to C transition was observed in the AT track of CArG_2 in *Capsella rubella, Capsella grandiflora* and *Eutrema salsugineum* (Figure 4J). Therefore, we tested whether AG, SEP3, and/or the SHP proteins could bind to these diverged CArG_2 motifs. We chose the sequences from *C. rubella*, as the CArG_2 motif was identical to *C. grandiflora* (Figure 4H), and *E. salsugineum* (Figure 4I), and generated probes that encompassed both CArG motifs. We observed near identical results when these probes were incubated with the above proteins when compared to the *AtMUTE_i1* probe (Figure 4H-I). This suggests that the interaction between *MUTE*, AG, SEP3, and to a lesser extent SHP1, is conserved in the Brassicaceae.

### Stomatal development on gynoecial valves upon *AG* perturbation

We next asked whether the elevated transcription of *MUTE* and *FAMA* observed in the plants lacking complete *AG* activity was sufficient to modify stomatal formation on the gynoecial valves. Using SEM, we imaged the gynoecia and siliques of wild-type L-*er* and *ag-10* plants (Figure 5A-B, D-E, I-J). At stage 12, as described above, mid and late-stage stomatal lineage cells are largely absent from wild-type gynoecial valves. In contrast, stage 12 *ag-10* gynoecial valves have eight times the number of mid-stage stomatal lineage cells (*p*.*adj* = 0.0003, pairwise unpaired two-tailed t-tests) (Figure 5A-B, I, Supplemental Table 7). At stage 13, mid (*p*.*adj* = 0.02, pairwise unpaired two-tailed t-tests) and late-stage (*p*.*adj* < 10^−7^, pairwise unpaired two-tailed t-tests) stomatal lineage cells were much more abundant on *ag-10* gynoecial valves in comparison to wild-type (Figure 5D-E, J, Supplemental Table 7). Concomitantly, we observed a severe reduction in the number of early-stage stomatal lineage cells (*p*.*adj* = 10^−6^, pairwise unpaired two-tailed t-tests) (Figure 5D-E, J, Supplemental Table 7). We also observed a substantial increase in the number of late-stage stomatal lineage cells after perturbation of *AG* activity through the *AG-amiRNA* relative to uninduced plants (*p*.*adj* < 10^−5^, pairwise unpaired two-tailed t-tests) or similarly treated L-*er* wild-type plants (*p*.*adj* < 10^−5^, pairwise unpaired two-tailed t-tests) (Supplemental Figure 6A-D, Supplemental Table 9). It has been previously observed that the mutant of another MADS-domain transcription factor-coding gene, *FRUITFULL* (*FUL*), does not bear stomata on the silique (Gu *et al*., 1998). Notably, introgression of the *shp1-1 shp2-1* mutations into the strong *ful-1* mutant results in the recovery of stomata on siliques (Ferrándiz *et al*., 2000*b*). We asked whether introduction of the *ag-10* mutant into the *ful-1* background would also be sufficient to rescue the formation of stomata on siliques. We found stomatal formation was partially restored on *ag-10 ful-1* siliques although the stomata are not completely normal in appearance and are not distributed throughout the silique (Figure 5G-H). Mean expression of *SHP1* was not reduced (*p* = 0.87, two-tailed paired t-test) and *SHP2* was only mildly reduced 0.87-fold (*p* = 0.12, two-tailed pared t-test) in the *ag-10* background compared to L-*er* (Supplemental Figure 6E, Supplemental Table 3). Therefore, recovery of stomata on *ag-10 ful-1* siliques is probably independent of *SHP* activity. Together these data demonstrate that AG is required to suppress stomatal development during gynoecium development.

**Figure 6.**
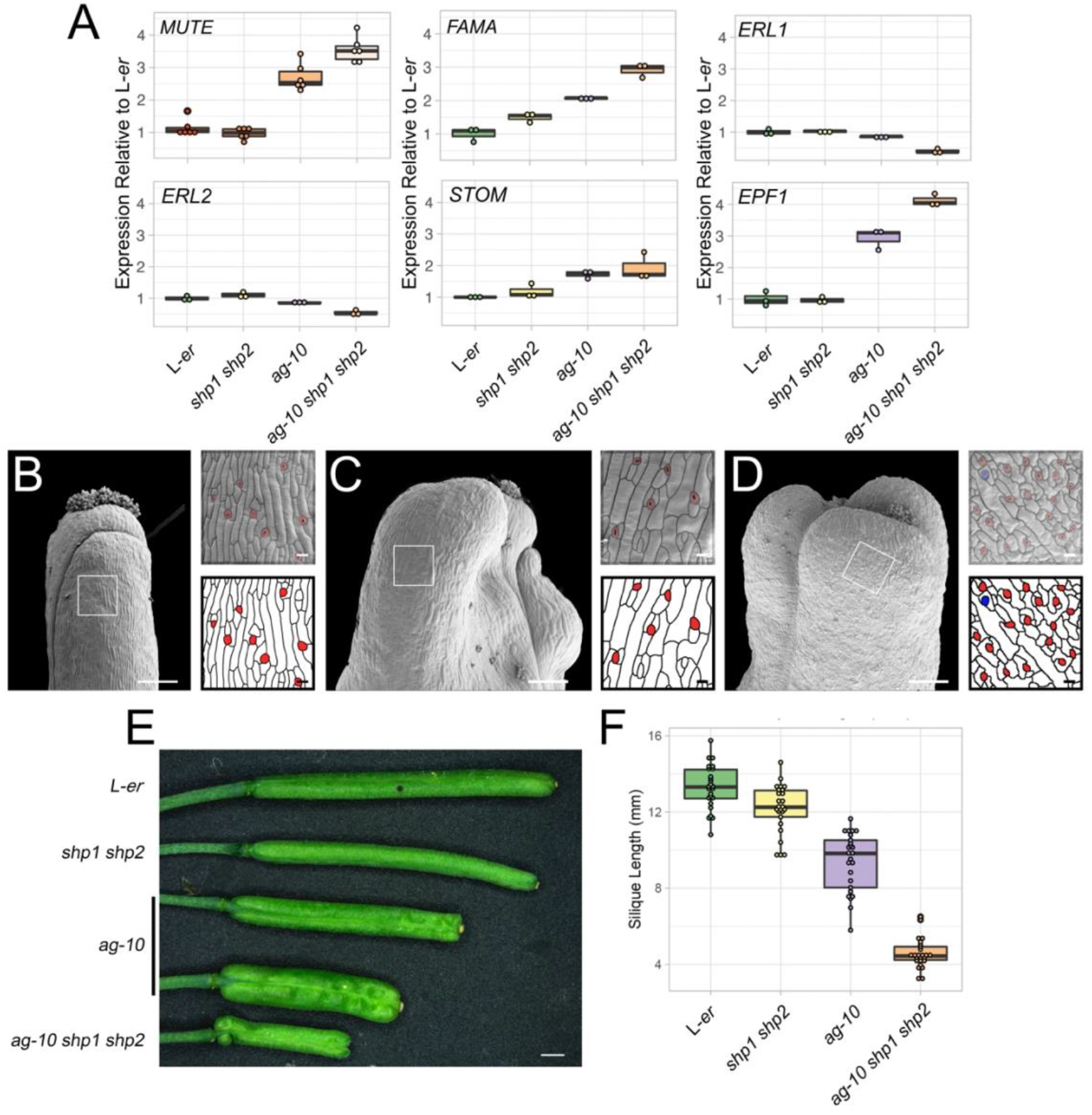
Redundancy between *AG* and *SHP1/2* during gynoecium and silique development. (A) Levels of *MUTE, FAMA, ERL1, ERL2, STOM*, and *EPF1* mRNAs as determined by RT-qPCR, in L-*er, shp1 shp2, ag-10, ag-10 shp1 shp2* stage 10-13 gynoecia relative to average of L-*er* samples. Each dot represents an individual independent biological replicate. (B-D) Scanning electron micrographs of mature siliques of (B) L-*er*, (C) *ag-10*, and (D) *ag-10 shp1 shp2*. Purple, blue, and red highlights indicate early, mid, and late-stage stomatal lineage morphology. Scale bars for images of whole gynoecia are 200 µm. Scale bars for magnifications are 20 µm. (E) Mature siliques of L-*er, shp1 shp2, ag-10, ag-10 shp1 shp2*. Scale is 1 mm. (F) Length of L-*er, shp1 shp2, ag-10, ag-10 shp1 shp2* mature siliques.

### Comparative activity of *AG* and *SHPs* during stomatal development

To understand more about the redundancy between *AG* and the *SHP* genes during stomatal development, we generated an *ag-10 shp1 shp2* triple mutant and examined the stomatal indices (i.e. the ratio cell belonging to the stomatal lineage to the total number of stomata and epidermal cells) on gynoecial and silique valves. Late-stage stomatal lineage cells were more common on the valves of triple mutant gynoecia at stage 12 relative to *ag-10* (*p*.*adj* = 0.03, pairwise two-tailed t-test). However, slightly fewer late-stage stomata formed on the valves of triple mutant gynoecia relative to *ag-10* at stage 13 (Figure 5B, C, E, F, I-J, Supplemental Table 7). We also tested the mRNA levels of *SPCH, MUTE*, and *FAMA* in this *ag-10 shp1 shp2* triple mutant background in comparison to parental genotypes (Figure 6A, Supplemental Figure 7E, Supplemental Table 3). Mean *SPCH* levels were reduced in the *ag-10 shp1 shp2* triple mutant relative to *ag-10* to ~0.84 fold, although the variation in expression was quite high (*p* = 0.55, pairwise two-tailed t-test) (Supplemental Figure 7E, Supplemental Table 3). Mean *MUTE* mRNA levels were elevated in the *ag-10 shp1 shp2* triple mutant relative to *ag-10* to ~1.33 fold (*p*.*adj* = 0.012, pairwise two-tailed t-test). *FAMA* mRNA levels were also elevated to ~1.4 fold in the *ag-10 shp1 shp2* triple mutant relative to *ag-10* (*p*.*adj* = 0.022, pairwise two-tailed t-test, Figure 6A, Supplemental Table 3). In contrast, levels of *SPCH, MUTE*, and *FAMA* mRNAs were not as strongly affected in the *shp1-1 shp2-1* double mutant background (Figure 6A, Supplemental Figure 7E, Supplemental Table 3). We concluded that the SHP proteins probably suppress the expression of *MUTE* in parallel with AG, which results in its elevated expression and the elevated expression of *FAMA* in the triple mutant background relative to *ag-10*.

Two other genes that are associated with AG binding are also involved in stomatal development, *ERL1* and *ERL2* (Ó’Maoiléidigh *et al*., 2013). These genes encode receptor-like kinases that control stomatal patterning in concert with TMM (Shpak *et al*., 2005). Ligands, such as STOMAGEN (STOM) and EPF1, bind to complexes of TMM, ER, ERL1, and ERL2, to maintain stomatal spacing and density (Torii, 2021). Therefore, we tested the mRNA levels of these genes in response to *AG* and *SHP* perturbation. *ERL1* and *ERL2* expression was mildly decreased in *ag-10*, however, their mRNA levels were reduced to ~0.4 (*p*.*adj* = 0.038, pairwise two-tailed t-test) and ~0.5-fold (*p*.*adj* = 0.009, pairwise two-tailed t-test) in the triple *ag-10 shp1 shp2* mutant relative to L-*er*, respectively (Figure 6A, Supplemental Table 3). *EPF1* mRNA levels were elevated in both *ag-10* and the triple mutant to ~3-fold (*p*.*adj* = 0.009, pairwise two-tailed t-test) and ~4.3-fold (*p*.*adj* = 0.009, pairwise two-tailed t-test), respectively (Figure 6A, Supplemental Table 3). *EPF1* levels were elevated by ~1.4-fold in the triple mutant relative to *ag-10* (*p*.*adj* = 0.03, pairwise two-tailed t-test) (Figure 6A, Supplemental Table 3). Similarly, mean *STOM* expression was elevated in *ag-10* and the triple mutant by ~1.7 (*p*.*adj* = 0.06, pairwise two-tailed t-test) and ~1.9-fold (*p*.*adj* = 0.14, pairwise two-tailed t-test) (Figure 6A, Supplemental Table 3). In contrast to these observations, mean *TMM* mRNA levels were not changed in *ag-10* or the *ag-10 shp1 shp2* triple mutant backgrounds (*p* = 0.39, two-way ANOVA) (Supplemental Figure 7E, Supplemental Table 3). Furthermore, there was little change in the expression of these genes in the *shp1 shp2* mutant background (Figure 6A, Supplemental Figure 7E-F, Supplemental Table 3).

Although the accumulation of *ERL1, ERL2, STOM*, and *EPF1* mRNAs were changed in the *ag-10* and/or the *ag-10 shp1 shp2* mutant backgrounds, we did not observe stomatal clustering in the samples in which *AG* or *SHP* were perturbed (Figure 6B-D). This suggests that either the observed changes are not substantial enough to modify patterning or that an initial change in gene expression mediated by *AG* and/or *SHP* was buffered by the regulatory network to maintain patterning. Notably, combinations of *er, erl1*, and *erl2* mutations results in a reduction of silique length (Shpak *et al*., 2004), which is similar to the reduction of silique length observed for *ag-10 shp1 shp2* plants relative to parental genotypes and wild-type (Figure 6E-F, Supplemental Table 10). Therefore, it is likely that AG-mediated control of *ERL1* and *ERL2* expression is related only to gynoecial growth rather than stomatal patterning.

## Discussion

The progression of stomatal development on the gynoecium/silique has not been previously described. Here, we show that stomatal development initiates on the gynoecial valves between late stage 11 and early stage 12 (Figure 2A-D, Supplemental Table 7). Stomatal initiation in the gynoecium correlates with the increased expression of stomatal bHLH transcription factors during this period (Figure 1). Most stomates do not reach maturity until after fertilization, although their maturation does not depend on fertilization (Supplemental Figure 2). This fertilization-independent differentiation is reminiscent of other differentiation processes, such as the continued increase in chlorophyll concentration in the gynoecia of emasculated pistils (i.e. unfertilized) (Carbonell-Bejerano *et al*., 2010), which also temporally correlate with but are not dependent on fertilization (Wagstaff *et al*., 2009). Given that fertilization does not control the onset or progression of stomatal development on gynoecial valves, we searched for an alternative mechanism.

The floral organ identity gene *AG* has been implicated previously in the differentiation of the gynoecial valve and in the suppression of leaf-like traits (Ó’Maoiléidigh *et al*., 2013, 2018). Here, we present evidence that AG forms a complex with SEP3 to suppress the formation of stomata (Figure 5), which are also typical of leaves, until the point of anthesis when their expression dissipates from gynoecial valves (Figure 2E-F, Supplemental Figure 3). AG and SEP3 accomplish this by directly repressing the transcription of *MUTE*, which encodes a key bHLH transcription factor required for stomatal development (Figure 3, 4). *MUTE* mRNA levels are elevated in the gynoecia of plants with reduced *AG* activity. Furthermore, the expression of *FAMA*, which is directly promoted by MUTE (Han *et al*., 2018), is elevated when *AG* activity is perturbed (Figure 3). By visualizing *FAMA* expression using a transgenic reporter line (Lee *et al*., 2019), we showed that perturbation of *AG* activity does not release *FAMA* expression throughout the epidermis of the gynoecium but only in what are likely stomatal lineage cells (Figure 3D-G). This suggests that SPCH drives *MUTE* expression in a similar manner as during leaf development and that AG counteracts this SPCH activity in stomatal lineage cells. *MUTE* expression gradually increases in stomatal lineage cells once AG protein levels are reduced after late stage 12, probably enabling SPCH to fully activate *MUTE* transcription (Figure 1). Therefore, stomatal lineage progression, but not entry, is modified by AG.

AG and SEP3 bind to the first intron of *MUTE*, which was previously demonstrated by ChIP-Seq and seq-DAP-seq experiments. The bound regions contain two CArG motifs and we showed that each CArG motif influences binding in a different way (Figure 4). We confirmed that both homodimers and heterodimers of AG and SEP3 can bind to the first intron of *MUTE*. We also showed that heterodimers of SHP1-SEP3 bind to the same region. Heterodimer binding is dependent on the presence of only one of these CArG motifs (CArG_2), whereas homodimers do not fully depend on the presence of CArG_2 although homodimer affinity is reduced in its absence. Mutagenesis of both CArG motifs, however, abolishes binding of both homodimers and heterodimers (Figure 4C-F). It has been suggested that AG-SEP3 quartets are required only to maintain floral meristem determinacy (Hugouvieux *et al*., 2018). Here, we did not observe the formation of higher order complexes with the generated probes suggesting that homodimers or heterodimers are sufficient to suppress *MUTE* expression. This is consistent with the idea that quartets of AG-SEP3 are not functional in the control of differentiation (Hugouvieux *et al*., 2018).

The observation that heterodimers of SHP1-SEP3 and, to a lesser extent, SHP2-SEP3 are also capable of binding to the *MUTE* first intron (Figure 4) is in keeping with observations of redundant activities between AG, SHP1, and SHP2 (Figure 5, Figure 6). At stage 12, stomatal lineage cells had progressed further in the triple mutant than *ag-10* while *shp1-1 shp2-1* mutants appeared similar to L-*er* (Figure 5). The progression of stomatal development on *ag-10* and the *ag-10 shp1 shp2* triple mutant was, however, similar at stage 13 although there was a large amount of variation associated with these measurements. Therefore, there is only weak phenotypic evidence to support the redundant control of stomatal development between *AG* and *SHP*. On a molecular level, there is stronger support for redundancy. Key regulatory genes, such as *MUTE* and *FAMA*, were differentially expressed to a greater degree in the triple mutant when compared to *ag-10* and were generally not differentially expressed in *shp1-1 shp2-1* double mutants relative to L-*er* (Figure 6). Patterning genes such as *ERL1, ERL2*, and *EPF1* show a similar trend although there is no change in stomatal clustering. As such, the effect of AG and SHP on *ERL1* and *ERL2* expression may be more related to gynoecium/silique growth than stomatal development. Considering both the molecular and phenotypic data, we conclude that functional redundancy between *AG* and *SHP* does play a role in the control of stomatal development and gynoecial/silique growth.

As previously suggested, the divergent expression patterns of *AG, SHP1*, and *SHP2* underlie many of their functional differences. The *SHPs* are broadly expressed in the gynoecium, including the valves, from stage 7 but become restricted to the valve margins during stage 12 of gynoecium development. This expression pattern is largely imposed by the FUL transcription factor, which represses *SHP* expression in the valves. Notably, *ful* mutants lack stomata on the valves although their formation can be restored when *SHP* activity is simultaneously perturbed. We observed a similar scenario as stomata were also recovered on *ag-10 ful-1* mutant valves (Figure 5G-H). This supports the idea that AG and the SHP proteins are suppressing stomatal development by repressing *MUTE* expression.

We propose that AG acts as a timer to control the emergence of stomata on the developing gynoecium/silique valve in *A. thaliana*. The conserved CArG motifs in the first intron of *MUTE* and the ability of AG, SEP3, and the SHPs to bind even slightly diverged sequences in *C. rubella* and *E. salsugineum* support the idea that this function is conserved in the Brassicaceae (Figure 4H-I). The correlation between stomatal development and fertilization suggests that atmospheric carbon assimilation by the silique stomata is used during silique photosynthesis to support seed development, particularly in terms of seed oil biosynthesis. Oils are stored in the seeds and mobilized during germination and early seedling development before autotrophic growth is established (Graham, 2008). Energy derived from silique photosynthesis supplements the accumulation of storage oils in the seeds in a variety of species including *A. thaliana* and *Brassica napus* (Sheoran *et al*., 1991; Gammelvind *et al*., 1996; Wang *et al*., 2016; Zhu *et al*., 2018).

The timing of stomatal formation on the silique valves may have been selected for a variety of reasons. Pre-anthesis formation of stomata may be disadvantageous due to selective pressure by pathogens (Wu and Liu, 2022). Pathogens can penetrate the plant through stomata and photosynthetic activity would probably be low in pre-anthesis gynoecia due to the shading effect of sepals. Furthermore, transpiration within a closed flower bud is unlikely to be particularly effective. These factors may have resulted in the selection for stomatal formation to coincide with flower bud opening. It is also possible that the timing of stomatal progression on gynoecial valves differs between species as stomatal development had not been completed even once siliques were exposed to light (e.g. stage 15-16 Supplemental Table 1). Therefore, there appears to be opportunity to initiate stomatal development at an earlier stage so that a higher number of active stomatal complexes are developed once the silique is exposed to light. If so, this may result from the modulation of AG function caused by divergence of the *cis* regulatory elements AG binds to. Assessing the correlation between stomatal development and fertilization in other species will partially address these questions.

## Acknowledgements

We thank Prof. Dominique Bergmann for her gift of seed lines (*SPCHpro:SPCH-YFP, FAMApro:NLS-2xYFP*, and *FAMApro:2xYFP*). We also thank Prof. Keiko Torii for her gift of the *MUTEpro:MUTE-GFP* seed line. We are grateful to Alison Beckett, Dr. Marco Marcello, and Dr. Marie Held for the technical assistance and to Dr. James Hartwell for use of equipment. We also thank Dr. Annette Becker and Clemens Rössner for providing us with functional data from their published work. DSOM was supported by a Humboldt Postdoctoral Fellowship and a BBSRC David Phillips Fellowship (BB/T009462/1). AJB is funded by the European Union’s Horizon 2020 research and innovation program under the Marie Sklodowska-Curie grant agreement No 897783. JM was funded by a BBSRC doctoral training partnership studentship. The laboratory of DSOM was funded by a BBSRC David Phillips Fellowship (BB/T009462/1). GC was funded by the Max Planck Society, a grant from the Deutsche forschungsgemeinschaft (https://www.dfg.de/, CO 318/11-1), a grant from the ERC (https://erc.europa.eu/, Nº339113 – HyLife) and is a member of a DFG-funded Cluster of Excellence (https://www.dfg.de/, EXC 2048/1 Project ID: 390686111).

## Notes

### Competing Interest Statement

The authors have declared no competing interest.

## References

Adrian J, Chang J, Ballenger CE, Bargmann Borr, Alassimone J, Davies KA, Lau OS, Matos JL, Hachez C, Lanctot A, Vatén A, Birnbaum KD, Bergmann DC. 2015. Transcriptome Dynamics of the Stomatal Lineage: Birth, Amplification, and Termination of a Self-Renewing Population. Developmental Cell 33, 107–118.

Alvarez J, Smyth DR. 2002. CRABS CLAW and SPATULA genes regulate growth and pattern formation during gynoecium development in Arabidopsis thaliana. International Journal of Plant Sciences 163, 17–41.

Bailey TL, Johnson J, Grant CE, Noble WS. 2015. The MEME Suite. Nucleic Acids Research 43, W39–W49.

Benjamini Y, Hochberg Y. 1995. Controlling the False Discovery Rate: A Practical and Powerful Approach to Multiple Testing. Journal of the Royal Statistical Society: Series B (Methodological) 57, 289–300.

Berardini TZ, Reiser L, Li D, Mezheritsky Y, Muller R, Strait E, Huala E. 2015. The arabidopsis information resource: Making and mining the ‘gold standard’ annotated reference plant genome. Genesis 53, 474–485.

Bowman JL, Drews GN, Meyerowitz EM. 1991a. Expression of the Arabidopsis floral homeotic gene AGAMOUS is restricted to specific cell types late in flower development. The Plant Cell 3, 749–758.

Bowman JL, Smyth DR, Meyerowitz EM. 1989. Genes directing flower development in Arabidopsis. The Plant Cell 1, 37–52.

Bowman JL, Smyth DR, Meyerowitz EM. 1991b. Genetic interactions among floral homeotic genes of Arabidopsis. Development 112, 1–20.

Carbonell-Bejerano P, Urbez C, Carbonell J, Granell A, Perez-Amador MA. 2010. A Fertilization-Independent Developmental Program Triggers Partial Fruit Development and Senescence Processes in Pistils of Arabidopsis. Plant Physiology 154, 163–172.

Cramer MD, Hawkins HJ, Verboom GA. 2009. The importance of nutritional regulation of plant water flux. Oecologia 161, 15–24.

Davies KA, Bergmann DC. 2014. Functional specialization of stomatal bHLHs through modification of DNA-binding and phosphoregulation potential. Proceedings of the National Academy of Sciences of the United States of America 111, 15585–15590.

Ditta G, Pinyopich A, Robles P, Pelaz S, Yanofsky MF. 2004. The SEP4 Gene of Arabidopsis thaliana Functions in Floral Organ and Meristem Identity. Current Biology 14, 1935–1940.

Dong J, Bergmann DC. 2010. Stomatal patterning and development. Current Topics in Developmental Biology 91, 267–297.

Edwards K, Johnstone C, Thompson C. 1991. A simple and rapid method for the preparation of plant genomic DNA for PCR analysis. Nucleic Acids Research 19, 1349.

Egea-Cortines M, Saedler H, Sommer H. 1999. Ternary complex formation between the MADS-box proteins SQUAMOSA, DEFICIENS and GLOBOSA is involved in the control of floral architecture in Antirrhinum majus. EMBO Journal 18, 5370–5379.

Favaro R, Pinyopich A, Battaglia R, Kooiker M, Borghi L, Ditta G, Yanofsky MF, Kater MM, Colombo L. 2003. MADS-Box Protein Complexes Control Carpel and Ovule Development in Arabidopsis. The Plant Cell 15, 2603–2611.

Ferrándiz C, Gu Q, Martienssen R, Yanofsky MF. 2000a. Redundant regulation of meristem identity and plant architecture by FRUITFULL, APETALA1 and CAULIFLOWER. Development 127, 725–734.

Ferrándiz C, Liljegren SJ, Yanofsky MF. 2000b. Negative regulation of the SHATTERPROOF genes by FRUITFULL during Arabidopsis fruit development. Science 289, 436–438.

Frazer KA, Pachter L, Poliakov A, Rubin EM, Dubchak I. 2004. VISTA: Computational tools for comparative genomics. Nucleic Acids Research 32, 273–279.

Fu LY, Zhu T, Zhou X, Yu R, He Z, Zhang P, Wu Z, Chen M, Kaufmann K, Chen D. 2022. ChIP-Hub provides an integrative platform for exploring plant regulome. Nature Communications 13.

Gammelvind LH, Schjoerring JK, Mogensen VO, Jensen CR, Bock JGH. 1996. Photosynthesis in leaves and siliques of winter oilseed rape (Brassica napus L.). Plant and Soil 186, 227–236.

Geisler M, Yang M, Sack FD. 1998. Divergent regulation of stomatal initiation and patterning in organ and suborgan regions of the Arabidopsis mutants too many mouths and four lips. Planta 205, 522–530.

Goethe J. 1790. Versuch die Metamorphose der Pflanzen zu erklären. Gotha, Germany: Ettinger.

Goodstein DM, Shu S, Howson R, Neupane R, Hayes RD, Fazo J, Mitros T, Dirks W, Hellsten U, Putnam N, Rokhsar DS. 2012. Phytozome: A comparative platform for green plant genomics. Nucleic Acids Research 40, 1178–1186.

Graham IA. 2008. Seed storage oil mobilization. Annual Review of Plant Biology 59, 115–142.

Gu Q, Ferrándiz C, Yanofsky MF, Martienssen R. 1998. The FRUITFULL MADS-box gene mediates cell differentiation during Arabidopsis fruit development. Development 125, 1509–1517.

Han SK, Qi X, Sugihara K, Dang JH, Endo TA, Miller KL, Kim ED, Miura T, Torii KU. 2018. MUTE Directly Orchestrates Cell-State Switch and the Single Symmetric Division to Create Stomata. Developmental Cell 45, 303–315.

Hara K, Kajita R, Torii KU, Bergmann DC, Kakimoto T. 2007. The secretory peptide gene EPF1 enforces the stomatal one-cell-spacing rule. Genes and Development 21, 1720–1725.

Honma T, Goto K. 2001. Complexes of MADS-box proteins are sufficient to convert leaves into floral organs. Nature 409, 525–529.

Hua W, Li RJ, Zhan GM, Liu J, Li J, Wang XF, Liu GH, Wang HZ. 2012. Maternal control of seed oil content in Brassica napus: The role of silique wall photosynthesis. The Plant Journal 69, 432–444.

Hugouvieux V, Jourdain A, Stigliani A, Carles CC, Parcy F, Charras Q, Zubieta C, Silva CS, Conn SJ, Conn V. 2018. Tetramerization of MADS family transcription factors SEPALLATA3 and AGAMOUS is required for floral meristem determinacy in Arabidopsis. Nucleic Acids Research 46, 4966–4977.

Hunt L, Gray JE. 2009. The Signaling Peptide EPF2 Controls Asymmetric Cell Divisions during Stomatal Development. Current Biology 19, 864–869.

Immink RGH, Tonaco IAN, de Folter S, Shchennikova A, van Dijk Adj, Busscher-Lange J, Borst JW, Angenent GC. 2009. SEPALLATA3: The ‘glue’ for MADS box transcription factor complex formation. Genome Biology 10.

Ito T, Ng KH, Lim TS, Yu H, Meyerowitz EM. 2007. The homeotic protein AGAMOUS controls late stamen development by regulating a jasmonate biosynthetic gene in Arabidopsis. The Plant Cell 19, 3516–3529.

Ito T, Wellmer F, Yu H, Das P, Ito H, Alves-Ferreira M, Riechmann JL, Meyerowitz EM. 2004. The homeotic protein AGAMOUS controls microsporogeniesis by regulation of SPOROCYTELESS. Nature 430, 356–360.

Kanaoka MM, Pillitteri LJ, Fujii H, Yoshida Y, Bogenschutz NL, Takabayashi J, Zhu JK, Torii KU. 2008. SCREAM/ICE1 and SCREAM2 specify three cell-state transitional steps leading to Arabidopsis stomatal differentiation. The Plant Cell 20, 1775–1785.

Kivivirta KI, Herbert D, Roessner C, de Folter S, Marsch-Martinez N, Becker A. 2020. Transcriptome analysis of gynoecium morphogenesis uncovers the chronology of gene regulatory network activity. Plant Physiology, 1–15.

Kondo T, Kajita R, Miyazaki A, Hokoyama M, Nakamura-Miura T, Mizuno S, Masuda Y, Irie K, Tanaka Y, Takada S, Kakimoto T, Sakagami Y. 2010. Stomatal density is controlled by a mesophyll-derived signaling molecule. Plant and Cell Physiology 51, 1–8.

Lai X, Stigliani A, Lucas J, Hugouvieux V, Parcy F, Zubieta C. 2020. Genome-wide binding of SEPALLATA3 and AGAMOUS complexes determined by sequential DNA-affinity purification sequencing. Nucleic Acids Research 48, 9637–9648.

Laux T, Mayer KFX, Berger J, Jürgens G, Genetik L, München L. 1996. The WUSCHEL gene is required for shoot and floral meristem integrity in Arabidopsis. Development 122, 87–96.

Lee LR, Wengier DL, Bergmann DC. 2019. Cell-type–specific transcriptome and histone modification dynamics during cellular reprogramming in the Arabidopsis stomatal lineage. Proceedings of the National Academy of Sciences of the United States of America 116, 21914–21924.

Liu X, Kim YJ, Müller R, Yumul RE, Liu C, Pan Y, Cao X, Goodrich J, Chen X. 2011. AGAMOUS Terminates Floral Stem Cell Maintenance in Arabidopsis by Directly Repressing WUSCHEL through Recruitment of Polycomb Group Proteins. The Plant Cell 23, 3654–3670.

Livak KJ, Schmittgen TD. 2001. Analysis of relative gene expression data using real-time quantitative PCR and the 2-ΔΔCT method. Methods 25, 402–408.

MacAlister CA, Ohashi-Ito K, Bergmann DC. 2007. Transcription factor control of asymmetric cell divisions that establish the stomatal lineage. Nature 445, 537.

Machanick P, Bailey TL. 2011. MEME-ChIP: Motif analysis of large DNA datasets. Bioinformatics 27, 1696–1697.

Mandel MA, Yanofsky MF. 1998. The Arabidopsis AGL9 MADS box gene is expressed in young flower primordia. Sexual Plant Reproduction 11, 22–28.

Melzer R, Verelst W, Theißen G. 2009. The class E floral homeotic protein SEPALLATA3 is sufficient to loop DNA in ‘floral quartet’-like complexes in vitro. Nucleic Acids Research 37, 144–157.

Nadeau JA, Sack FD. 2002a. Stomatal Development in Arabidopsis. The Arabidopsis Book 1, e0066.

Nadeau JA, Sack FD. 2002b. Control of stomatal distribution on the Arabidopsis leaf surface. Science 296, 1697–1700.

O’Maoileidigh DS, Graciet E, Wellmer F. 2014a. Gene networks controlling Arabidopsis thaliana flower development. New Phytologist 201, 16–30.

O’Maoileidigh DS, Graciet E, Wellmer F. 2014b. Genetic Control of Arabidopsis Flower Development. Advances in Botanical Research. Elsevier, 159–190.

Ó’Maoiléidigh DS, Stewart D, Zheng B, Coupland G, Wellmer F. 2018. Floral homeotic proteins modulate the genetic program for leaf development to suppress trichome formation in flowers. Development 145, dev157784.

Ó’Maoiléidigh DS, Wuest SE, Rae L, Raganelli A, Ryan PT, Kwasniewska K, Das P, Lohan AJ, Loftus B, Graciet E, Wellmer F. 2013. Control of Reproductive Floral Organ Identity Specification in Arabidopsis by the C Function Regulator AGAMOUS. The Plant Cell 25, 2482–2503.

Ohashi-Ito K, Bergmann DC. 2006. Arabidopsis FAMA Controls the Final Proliferation/Differentiation Switch during Stomatal Development. The Plant Cell 18, 2493–2505.

Pajoro A, Angenent GC, Kaufmann K, Madrigal P, Krajewski P, Muiño JM, Matus JT, Jin J, Riechmann J-L, Mecchia MA, Debernardi JM, Palatnik JF, Balazadeh S, Arif M, Ó’Maoiléidigh DS, Wellmer F, Krajewski P, Riechmann J-L, Angenent GC, Kaufmann K. 2014. Dynamics of chromatin accessibility and gene regulation by MADS-domain transcription factors in flower development. Genome Biology 15, R41.

Pasha A, Shabari S, Cleary A, Chen X, Berardini T, Farmer A, Town C, Provart N. 2020. Araport lives: An updated framework for Arabidopsis bioinformatics. The Plant Cell 32, 2683–2686.

Pelaz S, Ditta GS, Baumann E, Wisman E, Yanofsky MF. 2000a. B and C floral organ identity functions require SEPALLATA MADS-box genes. Nature 405, 200–203.

Pelaz S, Ditta GS, Baumann E, Wisman E, Yanofsky MF. 2000b. B and C floral organ identity functions require SEPALLATTA MADS-box genes. Nature 405, 200– 203.

Pelaz S, Tapia-Lopez R, Alvarez‐Buylla ER, Yanofsky MF. 2001. Conversion of leaves into petals in Arabidopsis. Current Biology 11, 182–184.

Pillitteri LJ, Sloan DB, Bogenschutz NL, Torii KU. 2007. Termination of asymmetric cell division and differentiation of stomata. Nature 445, 501.

Pyke KA, Page AM. 1998. Plastid Ontogeny during Petal Development in Arabidopsis. Plant Physiology 116, 797–803.

Ran JH, Shen TT, Liu WJ, Wang XQ. 2013. Evolution of the bHLH genes involved in stomatal development: Implications for the expansion of developmental complexity of stomata in land plants. PLoS ONE 8.

Ryan PT, Ó’Maoiléidigh DS, Drost HG, Kwaśniewska K, Gabel A, Grosse I, Graciet E, Quint M, Wellmer F. 2015. Patterns of gene expression during Arabidopsis flower development from the time of initiation to maturation. BMC Genomics 16, 1–12.

Sablowski R. 2015. Control of patterning, growth, and differentiation by floral organ identity genes. Journal of Experimental Botany 66, 1065–1073.

Sheoran I, Sawhney V, Babbar S, Singh R. 1991. In Vivo fixation of CO2 by attached pods of Brassica campestris L. Annals of Botany 67, 425–433.

Shpak ED, Berthiaume CT, Hill EJ, Torii KU. 2004. Synergistic interaction of three ERECTA-family receptor-like kinases controls Arabidopsis organ growth and flower development by promoting cell proliferation. Development 131, 1491–1501.

Shpak ED, McAbee JM, Pillitteri LJ, Torii KU. 2005. Stomatal patterning and differentiation by synergistic interactions of receptor kinases. Science 309, 290–293.

Smyth DR, Bowman JL, Meyerowitz EM. 1990. Early flower development in Arabidopsis. The Plant Cell 2, 755–767.

Team Rs. 2022. RStudio: Integrated Development Environment for R.

Theißen G, Melzer R, Rümpler F. 2016. MADS-domain transcription factors and the floral quartet model of flower development: linking plant development and evolution. Development 143, 3259–3271.

Thomson B, Wellmer F. 2019. Molecular regulation of flower development. Current Topics in Developmental Biology. Elsevier Inc., 185–210.

Torii KU. 2021. Stomatal development in the context of epidermal tissues. Annals of Botany 128, 137–148.

Urbanus SL, de Folter S, Shchennikova A V, Kaufmann K, Immink RGH, Angenent GC. 2009. In planta localisation patterns of MADS domain proteins during floral development in Arabidopsis thaliana. BMC Plant Biology 9, 5.

Wagstaff C, Yang TJW, Stead AD, Buchanan-Wollaston V, Roberts JA. 2009. A molecular and structural characterization of senescing Arabidopsis siliques and comparison of transcriptional profiles with senescing petals and leaves. The Plant Journal 57, 690–705.

Wang C, Hai J, Yang J, Tian J, Chen W, Chen T, Luo H, Wang H. 2016. Influence of leaf and silique photosynthesis on seeds yield and seeds oil quality of oilseed rape (Brassica napus L.). European Journal of Agronomy 74, 112–118.

Wasserstein RL, Schirm AL, Lazar NA. 2019. Moving to a world beyond p 0.05′. The American Statistician 73, 1–19.

Wu J, Liu Y. 2022. Stomata–pathogen interactions: over a century of research. Trends in Plant Science 27, 964–967.

Wuest SE, O’Maoileidigh DS, Rae L, Kwasniewska K, Raganelli A, Hanczaryk K, Lohan AJ, Loftus B, Graciet E, Wellmer F. 2012. Molecular basis for the specification of floral organs by APETALA3 and PISTILLATA. Proceedings of the National Academy of Sciences of the United States of America 109, 13452–13457.

Zhu X, Zhang L, Kuang C, Guo Y, Huang C, Deng L, Sun X, Zhan G, Hu Z, Wang H, Hua W. 2018. Important photosynthetic contribution of silique wall to seed yield-related traits in Arabidopsis thaliana. Photosynthesis Research 137, 493–501.

