## Supplemental Figures for "AGAMOUS mediates timing of guard cell formation during gynoecium development"

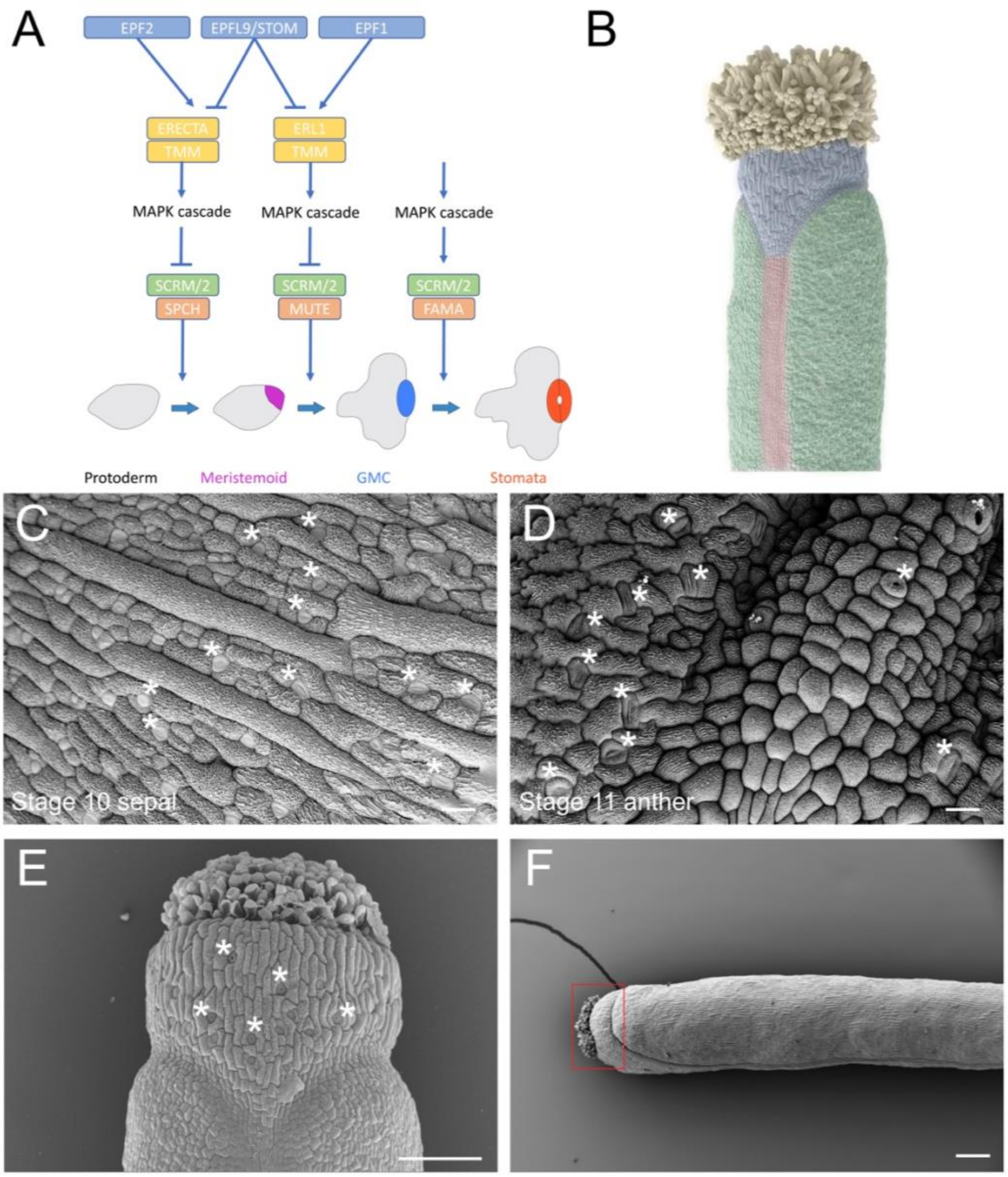


**Supplemental Figure 1. Overview of stomatal and gynoecium development and the presence of stomata on floral organs.** (A) Simplified model of stomatal development on leaves. The bHLH transcription factors, SPEECHLESS (SPCH), MUTE, FAMA, SCREAM (SCRM) and SCRM2 coordinate the progression of stomatal development. SPCH and MUTE are subject to post-translational regulation by a mitogen activated protein kinase (MAPK) cascade, which is activated by receptors, such as TOO MANY MOUTHS (TMM), ERECTA (ER) and ER-LIKEs (ERLs). The secreted peptides, EPIDERMAL PATTERNING FACTOR1 (EPF1) and EPF2 bind to these receptors to activate them at different stages of stomatal development. EPF-LIKE9/STOMAGEN (STOM) competes with EPF1/EPF2 binding to suppress activation of the receptors. (B) A scanning electron micrograph of a gynoecium at stage 13 (anthesis) with the valves (green), replum (red), style (blue) and stigma (yellow) false coloured. (C-E) Scanning electron micrographs of (C) stage 10 abaxial sepal, (D) stage 11 abaxial anther, (E) stage 12 gynoecium, and (F) stage >17 silique. Asterisks indicate presence of stomatal lineage cells. Red box in (F) highlights the size of style tissue in comparison to valve tissue, both of which bear stomata. Scale in (A-B) 20 μm, (C) 100 μm, and (D) 200 μm.

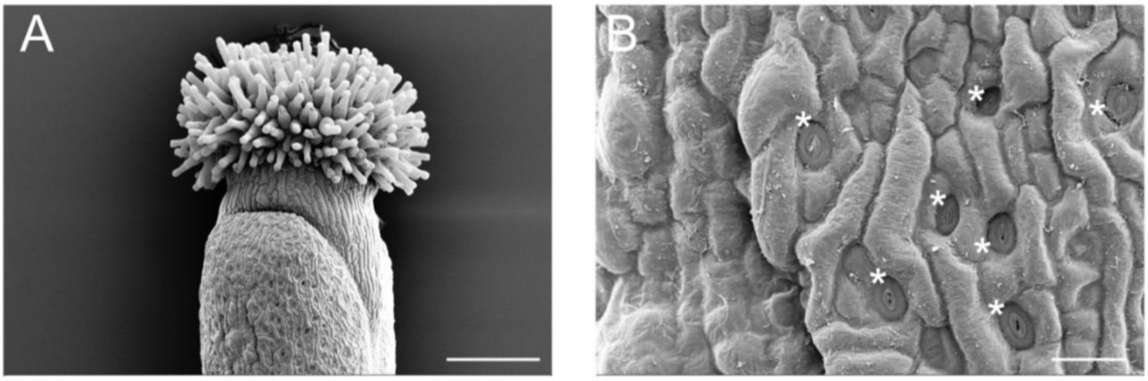


**Supplemental Figure 2. Presence of stomata on unfertilized gynoecial valve post-anthesis.** (A-B) Scanning electron micrographs of (A) a gynoecium from an emasculated flower 5 days after anthesis, (B) a magnification of the valve in (A). Scale in (A) 200 μm, (B) 20 μm.

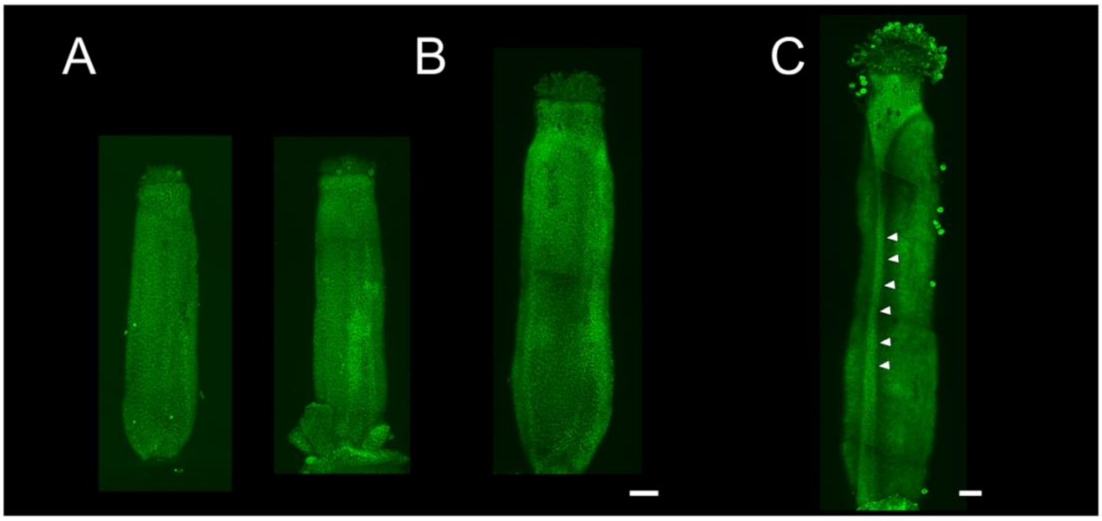


**Supplemental Figure 3. Confocal imaging of AG-GFP in late-stage gynoecia.** (A-C) Maximum intensity projections of stitched confocal laser scanning z-stack micrographs of (A) early stage 12, (B) late stage 12, and (C) stage 13 gynoecia from *AGpro:AG-GFP ag-1* plants. Arrowheads indicate accumulation of AG-GFP in the replum. Scale is 100 μm.

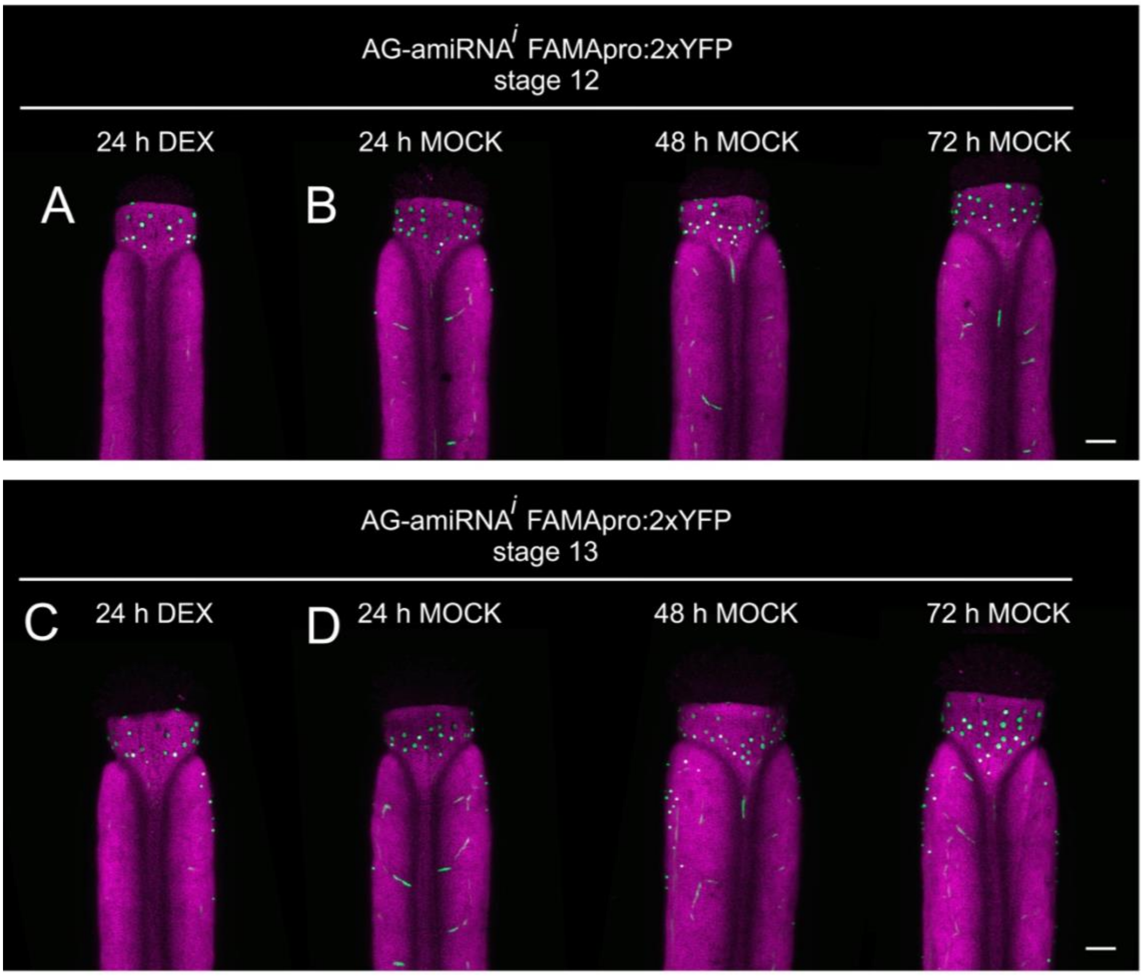


**Supplemental Figure 4. Confocal imaging of *FAMApro:2xYFP* in the *AG-amiRNA$^{i}$* background.** (A-D) Maximum intensity projections of stitched confocal laser scanning z-stack micrographs of *AG-amiRNA$^{i}$* (*OPpro:AG-amiRNA/35Spro:GR-LhG4*) *FAMApro:2xYFP* at (A-B) stage 12 and (C-D) stage 13 gynoecia after (A, C) dexamethasone or (B, D) mock treatments at the times indicated. Scale is 100 μm.

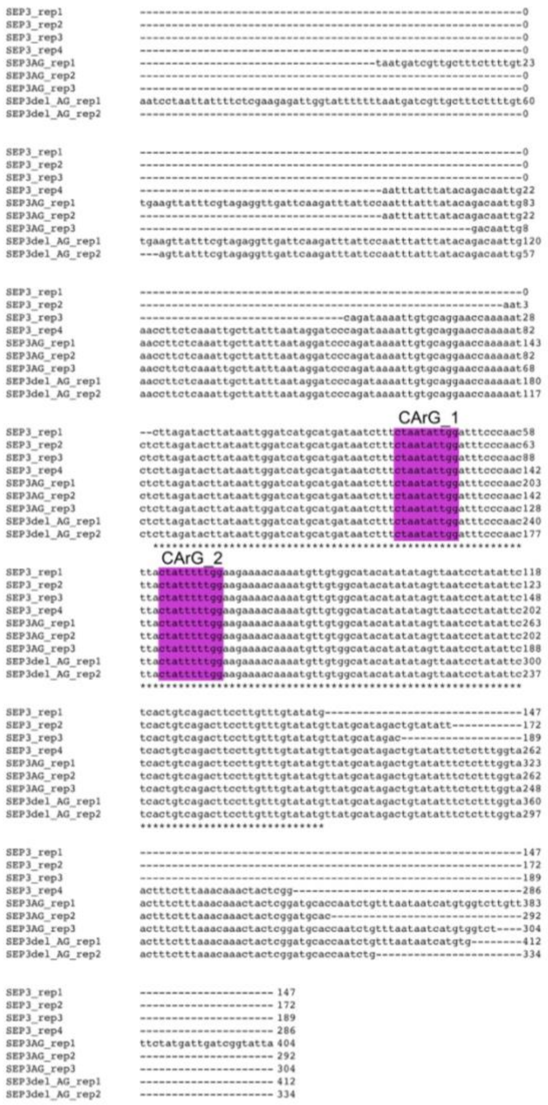


**Supplemental Figure 5. Sequences bound in *MUTE* first intron by AG-SEP3 from seq-DAP-seq.** Sequences mapping to the *MUTE* first intron identified as bound by SEP3 (SEP3_rep1-4), an AG-SEP3 complex (SEP3AG_rep1-3), or an AG-SEP3$^{\Delta tet}$ complex (SEP3del_AG_rep1-2) (Lai *et al.*, 2020). CArG_1 and CArG_2 are highlighted in purple boxes.

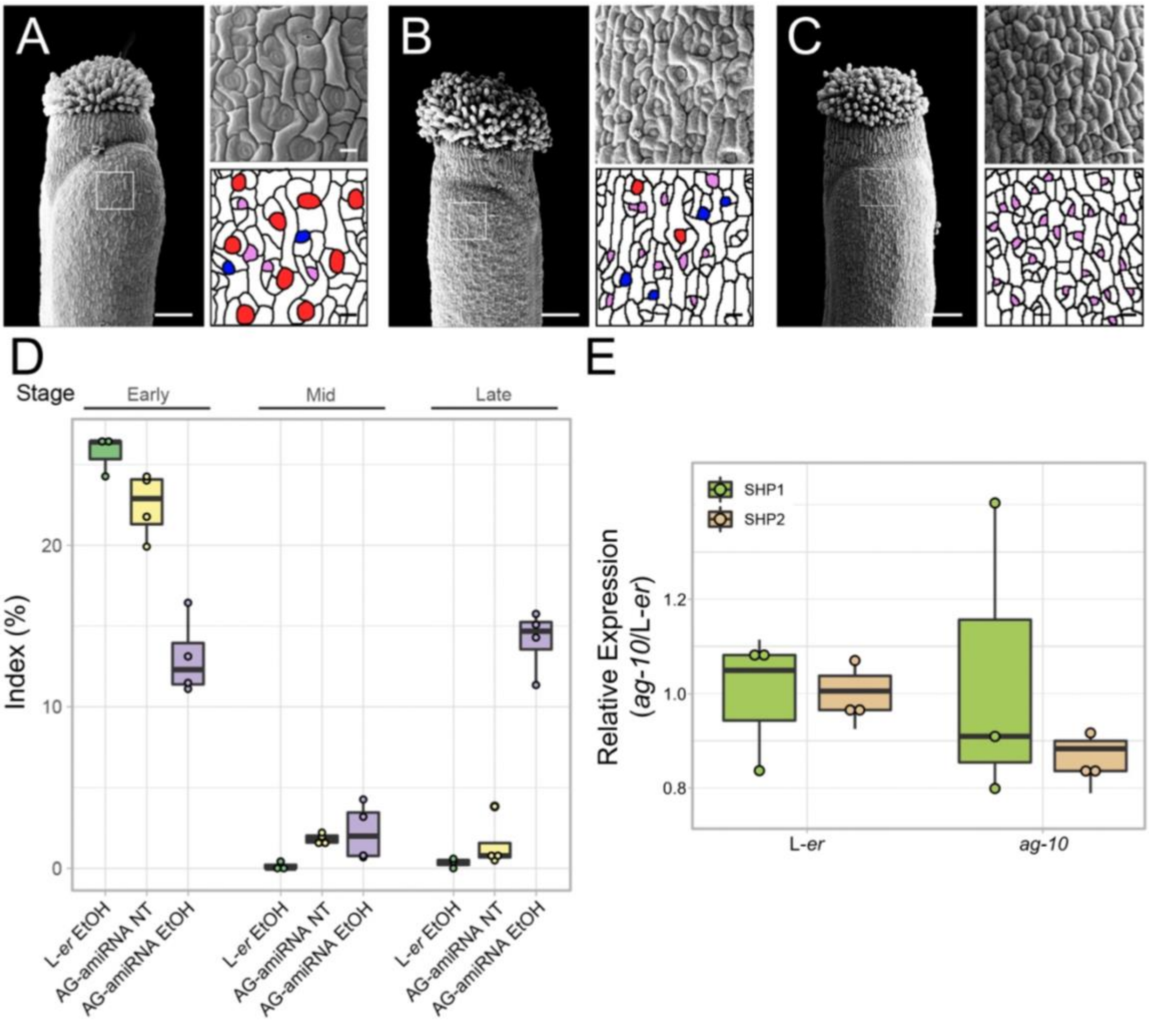


**Supplemental Figure 6. Stomatal development on gynoecial valves after *AG* knockdown.** (A-C) Scanning electron micrographs of gynoecia at anthesis of (A) *AlcRpro:AG-amiRNA/35Spro:AlcR* 5 d after 6 h EtOH vapour treatment, (B) untreated *AlcRpro:AG-amiRNA/35Spro:AlcR* and (C) L-*er* 5 d after 6 h EtOH vapour treatment. Scale bars for images of whole gynoecia are 100 µm. Scale bars for magnifications are 20 µm. Purple, blue, and red highlights indicate early, mid, and late-stage stomatal lineage morphology, respectively. (D) Index of early, mid, and late stomatal lineages based on morphology from scanning electron micrographs from stage 13 gynoecial valves of indicated genotypes. Each dot represents an individual sample. NT, no treatment. (E) Expression of *SHP1* and *SHP2* as determined by RT-qPCR in L-*er* and *ag-10* stage 10-13 gynoecia. Each dot represents an individual biological replicate.

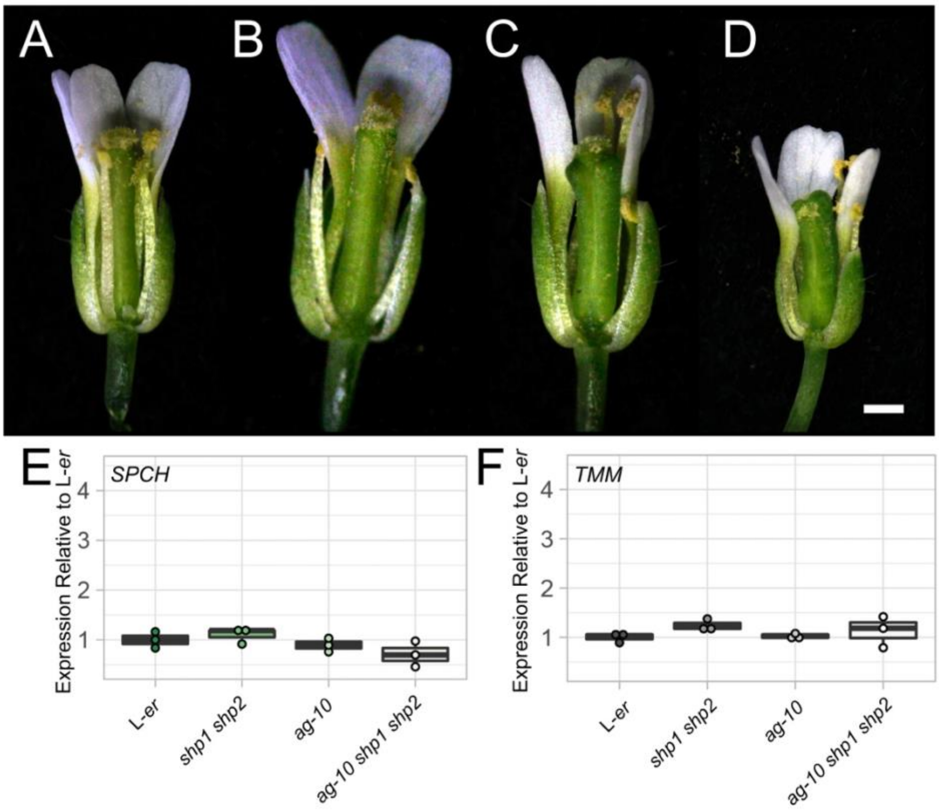


**Supplemental Figure 7. Morphology of flowers from mutant *AG* and *SHP* plants.** (A-D) Flowers at anthesis of (A) L-*er*, (B) *shp1-1 shp2-1*, (C) *ag-10*, and (D) *ag-10 shp1-1 shp2-1*. Some sepals, petals and stamens have been removed to allow visualization of gynoecium. Scale is 1 mm. (E-F) Levels of (E) *SPCH* and (F) *TMM* mRNAs in L-*er*, *shp1 shp2*, *ag-10*, *ag-10 shp1 shp2* stage 10-13 gynoecia as determined by RT-qPCR. Each dot represents an independent biological replicate.

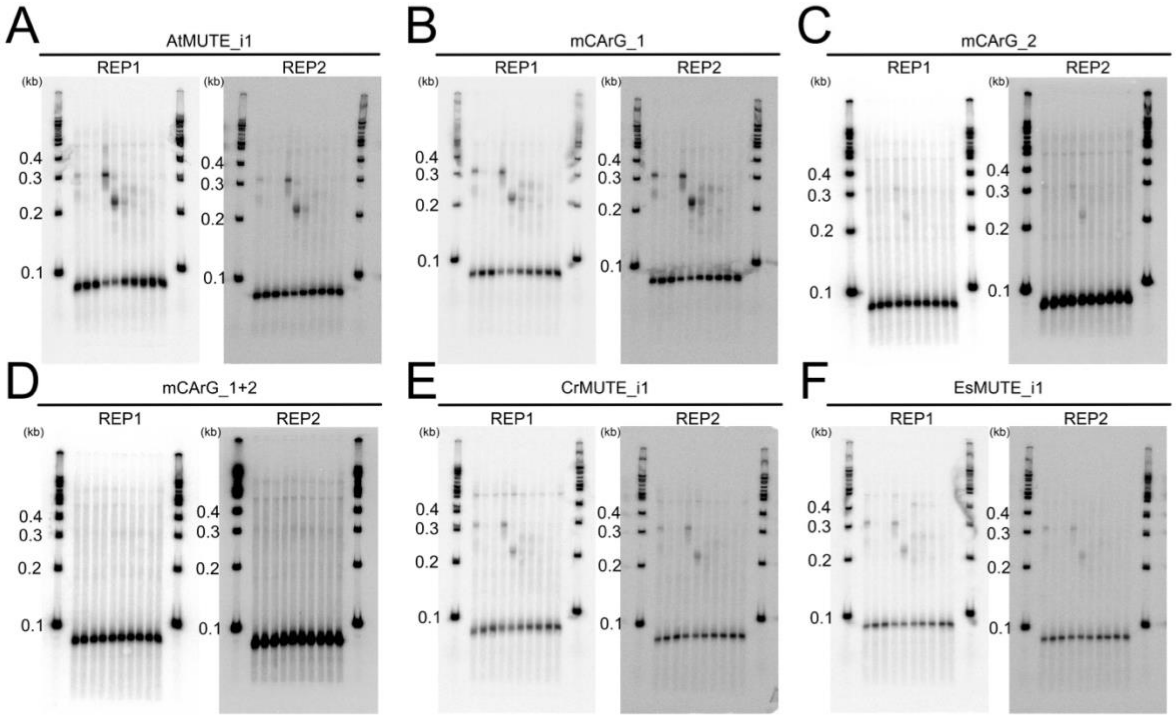


**Supplemental Figure 8. Gel shift assays of MADS domain transcription factors and the *MUTE* first intron.** (A-F) Protein-DNA gel shift assays using combinations of AG, SEP3, SEP3ΔC, SHP1, and SHP2 protein and two replicates of (A) *AtMUTE_i1*, (B) *mCArG_1*, (C) *mCArG_2*, (D) *mCArG_1+2*, (E) *CrMUTE_i1*, and (F) *EsMUTE_i1* probes.
